# Rapid PTEFb-dependent transcriptional reorganization underpins the glioma adaptive response to radiotherapy

**DOI:** 10.1101/2023.01.24.525424

**Authors:** Faye M. Walker, Lays Martin Sobral, Etienne Danis, Bridget Sanford, Ilango Balakrishnan, Dong Wang, Angela Pierce, Sana D. Karam, Natalie J. Serkova, Nicholas K. Foreman, Sujatha Venkataraman, Robin Dowell, Rajeev Vibhakar, Nathan A. Dahl

## Abstract

Dynamic regulation of gene expression is fundamental for cellular adaptation to exogenous stressors. PTEFb-mediated pause-release of RNA polymerase II (Pol II) is a conserved regulatory mechanism for synchronous transcriptional induction in response to heat shock, but this pro-survival role has not been examined in the applied context of cancer therapy. Using model systems of pediatric high-grade glioma, we show that rapid genome-wide reorganization of active chromatin facilitates PTEFb-mediated nascent transcriptional induction within hours of exposure to therapeutic ionizing radiation. Concurrent inhibition of PTEFb disrupts this chromatin reorganization and blunts transcriptional induction, abrogating key adaptive programs such as DNA damage repair and cell cycle regulation. This combination demonstrates a potent, synergistic therapeutic potential agnostic of glioma subtype, leading to a marked induction of tumor cell apoptosis and prolongation of xenograft survival. These studies reveal a central role for PTEFb underpinning the early adaptive response to radiotherapy, opening new avenues for combinatorial treatment in these lethal malignancies.

---

Adaptation and fine-tuning of gene expression are required to ensure appropriate cellular responses to developmental signals or environmental stressors. Neoplastic cells incorporate numerous signaling inputs in order to calibrate their net transcriptional output, including both cell state- and oncogene-specific intrinsic programs as well as extrinsic pressures from the tumor microenvironment, the immune system, and anti-cancer therapies. One of the most ubiquitously employed modalities in cancer is ionizing radiation (IR), in which genomic integrity is disrupted through generation of DNA double-strand breaks ^1^. The adaptive response to this cellular injury involves a coordinated interplay between DNA repair mechanisms and cell cycle regulation collectively termed the DNA damage response (DDR) ^1-3^. In cancer, effective DDR facilitates recovery from therapeutic IR and ultimately disease recurrence after treatment ^4-7^. This signaling network has been exhaustively well characterized, transduced through key regulatory proteins such ATM, ATR, and p53 ^2^. Much of this response occurs at a proteomic level, but robust transcriptomic changes are required to both initiate and sustain this corrective program ^8-10^. And underpinning the control of gene expression is an array of epigenetic inputs such as chromatin state and transcriptional cofactors which collectively regulate the recruitment, initiation, and productive elongation of RNA polymerase II (Pol II).

Focal chromatin state is the landscape over which transcription proceeds, including the physical accessibility of chromatin-binding factors to DNA as well as the net epigenetic signal conveyed by histone tail post-translational modifications (PTMs) ^11,12^. Many epigenetically active loci are relatively stable within a given cell, reflecting lineage commitment and developmental cell identity ^13,14^. But others are highly dynamic, reorganizing active chromatin to facilitate the transcriptional changes necessary to respond to a given stimulus ^15^. For example, fibroblasts have been shown to dramatically redistribute chromatin accessibility and H3K27ac over time following UV exposure, remodeling enhancers and super-enhancers in response to environmental pressure ^16^. Similar responses have been described across cell types and exogenous stimuli, highlighting the fundamental role of epigenetic signaling in driving adaptive transcriptional programs ^8,9,17,18^.

The functional endpoint of this epigenetic reorganization is productive transcriptional elongation by Pol II at the appropriate genomic loci. Processive transcription by Pol II is controlled through dynamic phosphorylation of the highly conserved C-terminal domain (CTD) ^19-21^. Following Pol II recruitment, initiation is governed by the pre-initiation complex, in which the CDK7-containing TFIIH phosphorylates the CTD at the Ser5 position ^22^. After transcribing 20-80 bases downstream, Pol II then becomes paused in promoter-proximal regions at many genomic loci. Phosphorylation of the CTD Ser2 by the CDK9/CyclinT1 pair, collectively termed positive transcription elongation factor b (PTEFb), is then required for pause release ^21,23,24^. This paused state can then represent a rate-limiting regulatory intermediate. Coordinated pause release is well defined in models of exogenous stressors such as heat shock ^25-27^ and developmental signaling ^28,29^, while misregulation of this checkpoint has been implicated in cancer and other human disease ^30-34^. Much has been learned about the mechanistic biology contributing to transcriptional elongation checkpoint control, but the therapeutic potential for its judicious disruption remains largely unrealized. Pediatric high-grade gliomas (HGGs), including diffuse intrinsic pontine gliomas (DIPGs) and other histone-mutant diffuse midline gliomas (DMGs), are aggressive malignancies of childhood for which radiation therapy remains the only standard of care, and HGG relapse after radiotherapy represents a uniformly fatal event. Here, we examine the dynamics of chromatin reorganization and PTEFb-mediated transcriptional induction in response to radiotherapy in pediatric HGG, and we investigate whether pharmacologic disruption of this adaptive reprogramming can be employed for therapeutic effect.

## Results

### Glioma cells rapidly reorganize active chromatin following exposure to IR

The physical accessibility of DNA determines permissible interactions by chromatin-binding factors to collectively regulate gene expression. The landscape of accessibility is rapidly and dynamically regulated in response to both extrinsic stimuli and developmental cues, and it can be measured by susceptibility of DNA to cleavage using techniques such as the assay for transposase-accessible chromatin using sequencing (ATAC-seq) (reviewed by Klemm et al ^11^). We first sought to characterize the early reorganization of accessibility that we hypothesized would occur immediately following radiotherapy. To do this, we utilized the SU-DIPG-IV (*HIST1H3B*^mut^, *ACVR1*^G328V^, *TP53*^WT^) culture model and performed ATAC-seq both at resting state and four hours following a single exposure to 6 Gy IR. This revealed a rapid shift biased directionally towards a more compacted chromatin state, with 3,237 peaks differentially lost and only 428 peaks differentially gained (**Figure 1a-b**). These accessibility losses occurred primarily at annotated promoters (79%, n=2,558), with only 21% (n=679) occurring at enhancers or other non-promoter transcription start sites (TSS). Ontology analysis of these loci showed functional enrichment in networks governing transcriptional processing and chromatin remodeling (e.g. *MED* subunits, *RBPJ, CHD4, EZH2*), DDR programs such as DNA repair and cell cycle regulation (*DUSP1, PPM1D, WEE1*), and cell structure morphogenesis (*NRCAM, STMN1, NPTN*) (**Figure 1c** and **Extended Data Table 1a**). These signatures suggest a selectivity for this chromatin compaction at loci functionally correlated with early adaptive cellular reorganization and DDR activation. Ontology analysis of differentially gained peaks yielded comparatively weak enrichment across nonspecific terms (**Extended Data Table 1b**). TRRUST analysis of transcriptional regulatory networks ^35,36^ within loci of differential accessibility inferred TP53 as the strongest upstream regulator affecting these changes, followed by HIF1A and E2F1 (**Figure 1d**). In order to better characterize active chromatin features which might predict changes in transcriptional output, we then performed ChIP-seq for H3K27ac, an epigenetic marker of transcriptionally active chromatin, in the same experimental conditions. In contrast to the largely unidirectional shift in accessibility, early H3K27ac redistribution was balanced, with 3,536 peaks differentially gained and 3,195 peaks differentially lost (**Figure 1e**). Peaks gained were almost exclusively at annotated promoters (94%, n=3,328), whereas losses were distributed across genomic elements (TSS 35%, n=1,132 vs eTSS 65%, n=2,063). When examined in relationship to the prior accessibility changes, the few regions of accessibility gains paradoxically lost acetylation, consistent with the nonspecific or secondary nature of this small subset of genomic elements. In contrast, the predominant cluster of promoters which underwent physical compaction after IR maintained comparatively stable levels of net acetylation, instead exhibiting a balanced redistribution of H3K27ac gains and losses across these regions (**Figure 1f**). Ontology analysis of differential H3K27ac peaks showed similar functional enrichments across networks mediating DDR programs, cell cycle and growth, and cell morphogenesis. Intriguingly, we observed a striking overlap in biological processes enriched in both H3K27ac gains and losses as opposed to distinct terms differentially up- or downregulated from these gene lists (**Figure 1g** and **Extended Data Table 2**). Taken together, these data reveal a rapid shift towards chromatin compaction within hours of exposure to therapeutic IR. This loss of accessibility is neither uniform nor random, but rather it is preferentially enriched in functional programs one would predict would be transcriptionally relevant in an early cellular DNA damage response. Within this shifting landscape of permissible physical interaction, we observe a broad redistribution of H3K27ac at gene promoters, creating a framework of active chromatin over which differential recruitment and activation of transcriptional machinery might now occur.

**Figure 1.**
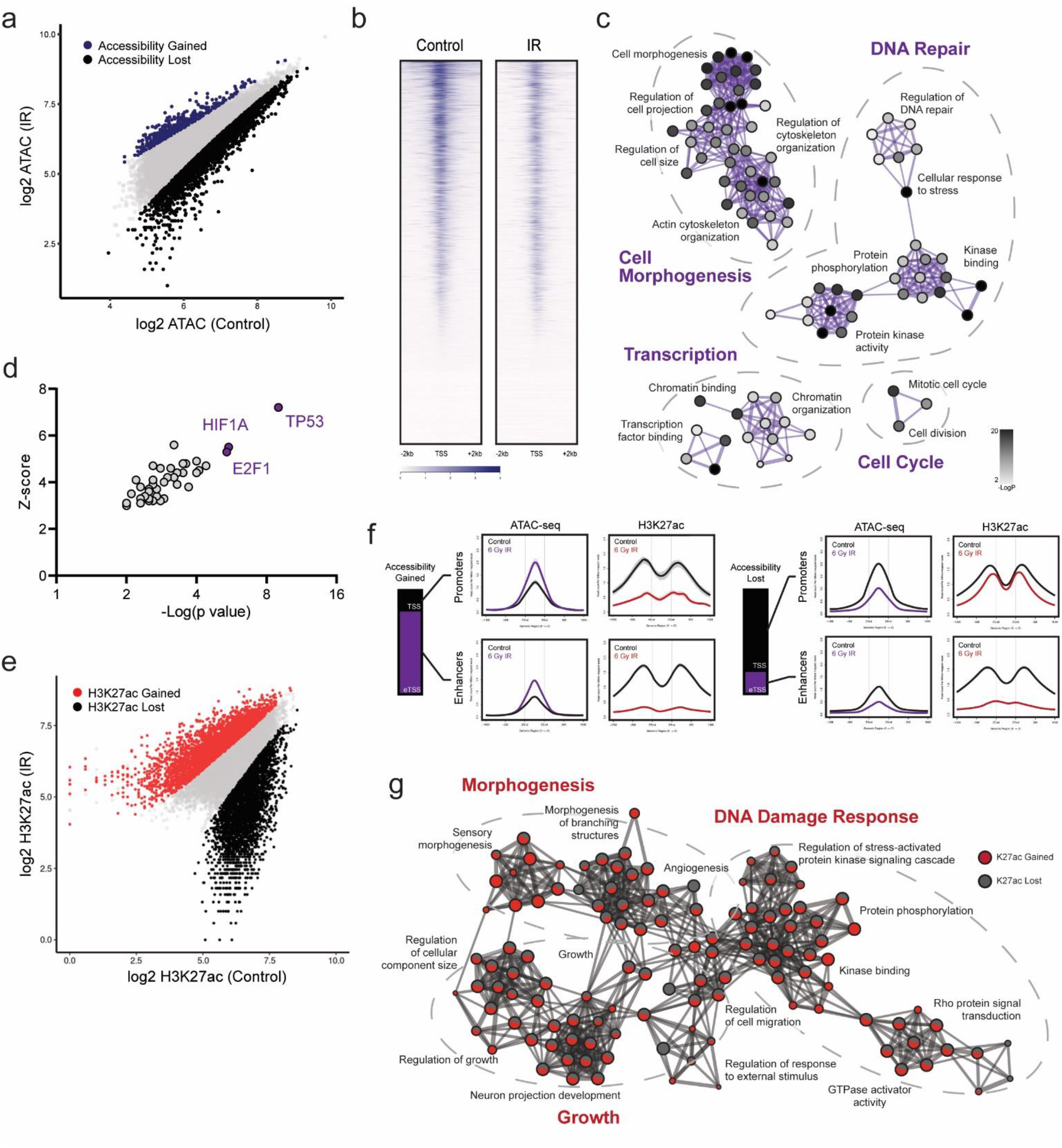
Glioma cells rapidly reorganize active chromatin following exposure to IR. **a**. Scatterplot of ATAC-seq peaks compared between IR-exposed cells and untreated controls (n=2). Differentially gained loci of accessibility are indicated in purple, differentially lost accessibility in black. **b**. Genome wide heatmap of accessibility before and after IR exposure. **c**. Gene ontology network constructed from loci of accessibility lost following IR. Each node denotes an enriched term, with color density reflecting - log(Pval). **d**. TRRUST inference of transcription factor-target pairs from differential ATAC-seq loci. **e**. Scatterplot of H3K27ac ChIP-seq peaks compared between IR-exposed cells and untreated controls (n=2). Differentially gained H3K27ac occupancy is indicated in red, differentially lost occupancy in black. **f**. H3K27ac ChIP-seq metagene profiles clustered by ATAC-seq-defined chromatin features. **g**. Gene ontology network constructed from loci of differential H3K27ac occupancy following IR. Each node denotes an enriched term, with color ratio reflecting relative contribution from H3K27ac gain and lost lists.

### Redistribution of H3K27ac occupancy correlates with early differential transcript expression

Prior work has demonstrated that DNA damaging events transiently decrease elongation rates, phenocopying slow Pol II mutants ^9,37^. Consistent with this, in situ fluorescent staining of nascent RNA synthesis showed a time-dependent decrease in net transcriptional output following a single IR exposure (**Figure 2a**). While this slowing enables repair of Pol II-encountered DNA lesions and limits mutagenesis, it is held in opposition with the need to rapidly activate transcriptional programs involved in early DDR ^8,9^. H3K27 acetylation functions as a transcriptionally activating modification in large part by serving as a target for transcription cofactor binding. This includes distinct PTEFb-containing protein complexes such as the bromodomain and extraterminal domain family protein BRD4 and the super elongation complex (SEC) ^38-41^. Once recruited in a catalytically active complex, the CDK9 subunit of PTEFb phosphorylates the CTD of Pol II at the serine 2 position, signaling for release into the gene body for productive elongation ^19-21,42^ (**Figure 2b**). Given the complex reordering of active chromatin we observed, we hypothesized that a similarly nuanced reorganization of transcriptional machinery might drive DDR programs within the context of overall transcriptional slowing.

**Figure 2.**
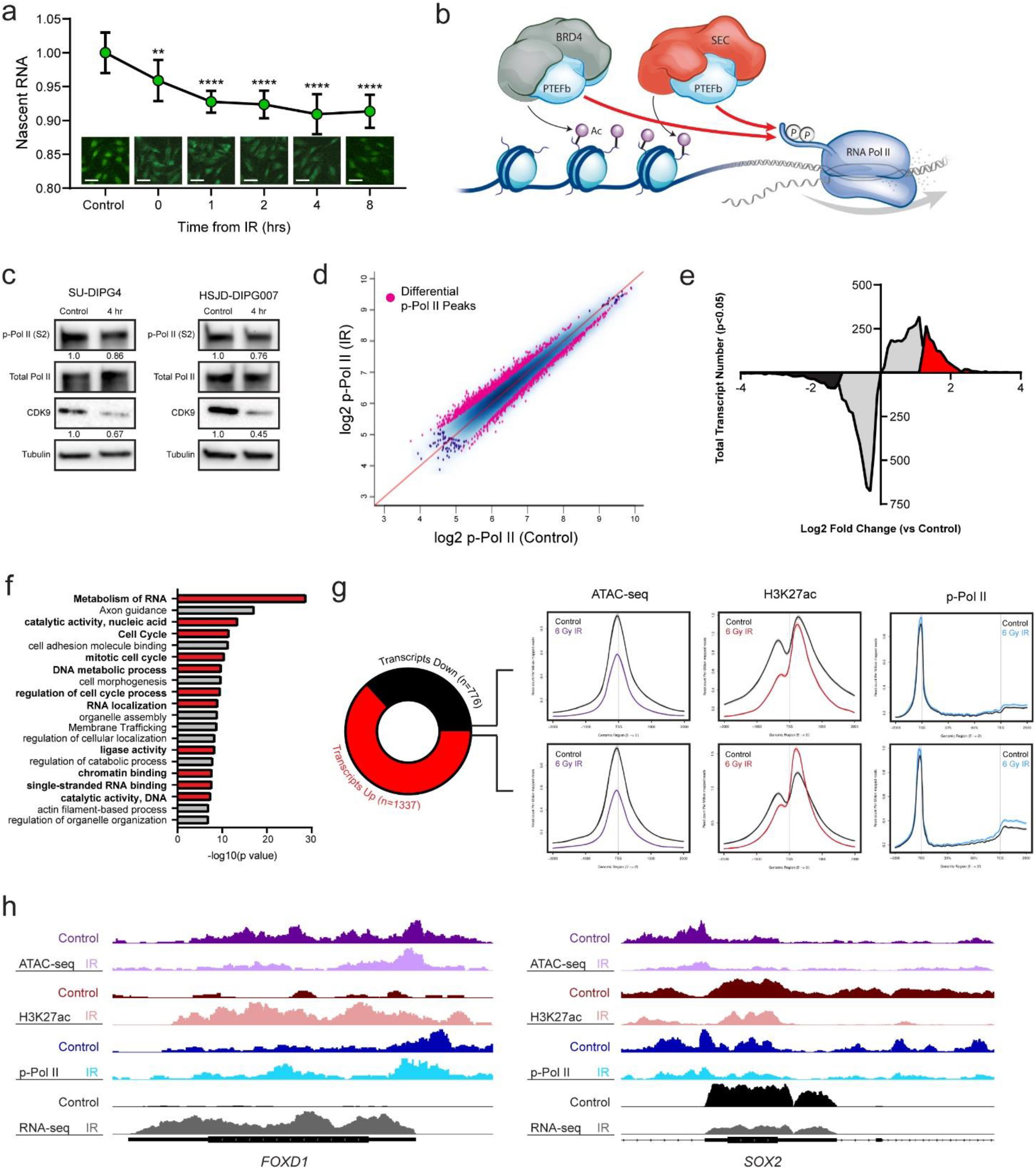
Redistributed H3K27ac occupancy correlates with early transcription from DDR programs. **a**. Click-IT fluorescent assay of relative nascent RNA abundance at indicated timepoints following IR. Comparisons reflect two-tailed Student’s t-test vs untreated control (** p=<0.01, **** p=<0.0001) (bar = 50 µm). **b**. Schematic representation of PTEFb localization to H3K27ac-marked chromatin by active BRD4- or SEC-PTEFb complexes to facilitate phosphorylation of Pol II CTD (Ser2). **c**. Immunoblot of p-Pol II (Ser2), total Pol II, and CDK9 measured 4 hours after 6 Gy IR. Value below represents mean quantification of biological triplicates. **d**. Scatterplot of p-Pol II (S2) CUT&RUN peaks compared between IR-exposed cells and untreated controls (n=3). Differentially bound peaks are indicated in pink. **e**. Histogram of differentially expressed transcripts following IR. Transcripts with significant (qval <0.05) but < 1.2 LF change are indicated in grey. Transcripts with > +/- LFC are in red and black, respectively. **f**. Functional ontology enrichment of transcripts ≥ 1.2 LFC in (e). Unbiased top 20 terms are displayed, with terms involved in transcriptional processing or DDR in red. **g**. Metagene plots of ATAC-seq, H3K27ac ChIP-seq, and p-Pol II (S2) CUT&RUN changes at differentially expressed transcripts. **h**. Illustrative loci at *FOXD1* and *SOX2* promoters demonstrate p-Pol II downstream egress and active transcription correlates with H3K27ac deposition irrespective of change in accessibility. Paired tracks reflect same data scale.

We first examined immunoblots of CDK9 expression and the CDK9-catalyzed Pol II (Ser2) phosphorylation mark at the same early timepoint as the above chromatin studies. These showed a modest decrease in total CDK9 expression and activity, respectively, concordant with net measures of transcriptional output (**Figure 2c**). In order to resolve occupancy profiles of CDK9 activity on chromatin, we next performed cleavage under targets and release using nuclease (CUT&RUN) for Ser2-phosphorylated Pol II. This revealed a striking redistribution p-Pol II (Ser2) activation across the genome within hours of IR exposure (1,923 reproducible, differential peaks p <0.05, n=3 replicates) (**Figure 2d** and **Extended Data Figure 1a-b**). Net change in chromatin occupancy was not significant using multiple peak calling parameters, suggesting that overall levels of Pol II activation are largely stable in this early time period (**Extended Data Figure 1c**). To define transcriptional output, we utilized transcriptome sequencing (RNA-seq) in the same conditions. This again demonstrated a greater number of unique transcripts differentially downregulated than up (4,775 down vs 3,677 up, qval < 0.05, n=3 replicates). In most cases, however, these downregulations were modest, falling below the 1.2 LFC cutoff commonly utilized for differential expression analyses. When examining only transcripts with differential LFC > +/- 1.2, this pattern was reversed, with a greater number of transcripts strongly induced than repressed (1,337 up vs 776 down, qval < 0.05) (**Figure 2e**). Gene ontology enrichment analysis of these upregulated transcripts revealed marked activation of programs involved in transcriptional processing, cell cycle regulation, and DNA catalytic activity, consistent with a focal induction of critical DDR programs despite global transcriptional slowing (**Figure 2f** and **Extended Data Table 3**).

We then intersected these RNA-seq data with the ATAC-seq, H3K27ac, and p-Pol II datasets. Surprisingly, differential expression by RNA-seq showed no correlation with changes in accessibility, as both up- and downregulated transcripts showed similar degrees of chromatin compaction following IR exposure (**Figure 2g**). This suggests that the primary function of this compaction is not in modulating gene expression, but it may instead reflect the protective role of a more heterochromatin state against subsequent DNA damage ^43,44^ for loci that must maintain active transcriptional output. In contrast, changes in H3K27ac deposition did exhibit correlation with differential transcript level, with an increase in the PTEFb-catalyzed p-Pol II pause-release and egress into downstream gene bodies observed at these loci (**Figure 2g-h**). In total, this demonstrates a central role for H3K27ac redistribution in organizing transcriptional output, including focal induction of critical DDR programs, within the broad chromatin compaction and transcriptional slowing observed following IR-induced genotoxic stress.

### PTEFb inhibition disrupts IR-induced chromatin reorganization and abrogates transcriptional induction

Given the role of H3K27ac in recruiting active PTEFb-containing complexes to drive Pol II pause-release, we hypothesized that selective inhibition of CDK9 should disrupt this adaptive response. To test this, we utilized AZD4573, a highly selective inhibitor of CDK9 with rapid target engagement, CDK9 enzymatic IC_50_ of <0.003 µmol/L, and >25-fold cellular selectivity for CDK9 versus other CDKs ^45^. Within our model system, we confirmed that AZD4573 treatment led to a dose-dependent depletion of both the CDK9-catalyzed Pol II Ser2 phosphorylation mark and nascent RNA synthesis (**Figure 3a-b**), supporting its utility for this perturbation.

**Figure 3.**
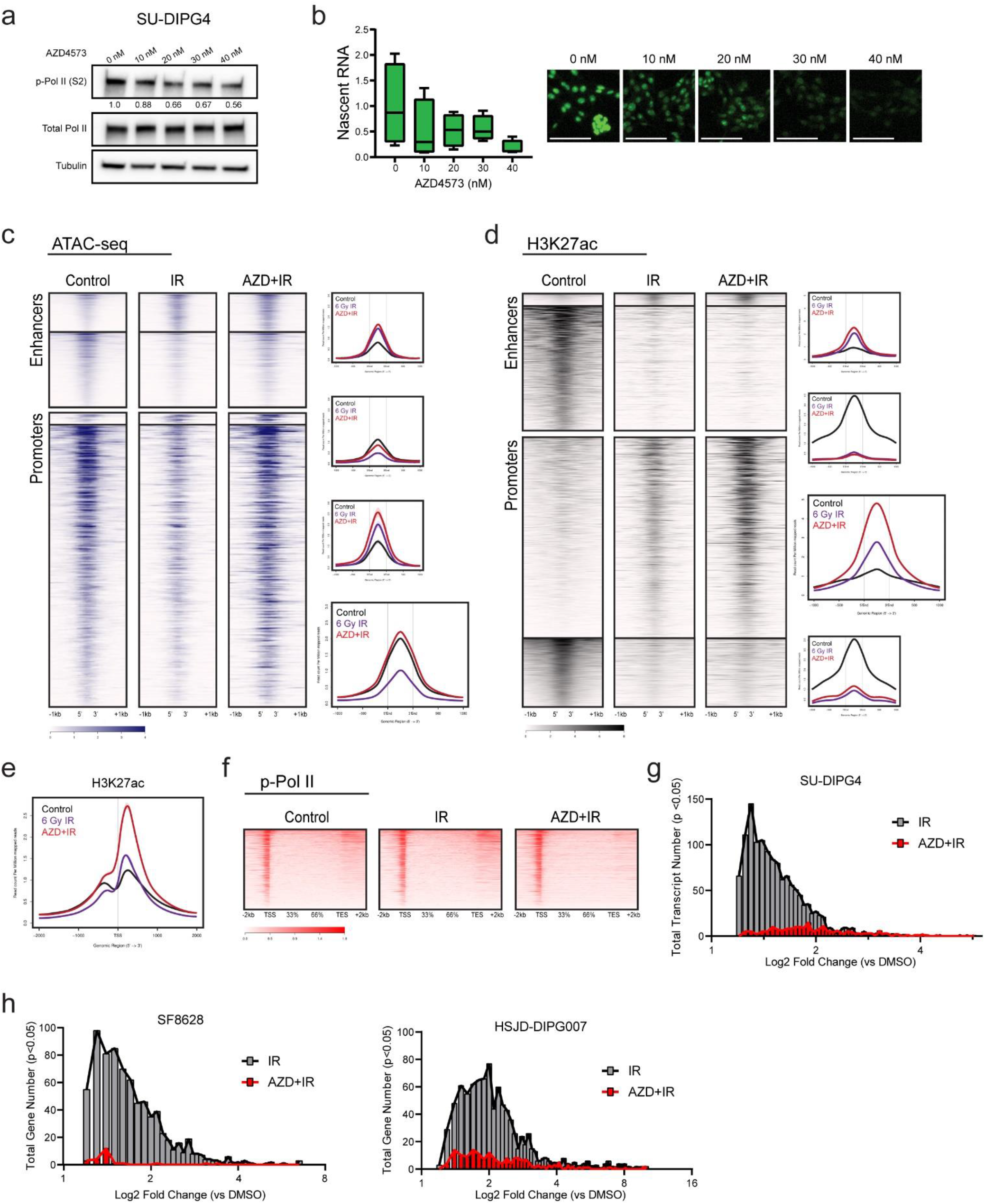
Concurrent CDK9 inhibition disrupts IR-driven chromatin reorganization and abrogates transcriptional induction. **a**. Immunoblot of p-Pol II (Ser2) and total Pol II at indicated doses of AZD4573. **b**. Click-IT fluorescent assay of relative nascent RNA abundance at indicated doses of AZD4573 (bar = 100 µm). **c**. ATAC-seq heatmap of untreated controls, IR exposed, or IR exposed with concurrent AZD4573 (40 nM) (n=2). Genome is clustered by change in ATAC-seq peaks following IR exposure. Metagene profile for each cluster is on right, with profile of the dominant cluster emphasized. **d**. H3K27ac heatmap of untreated controls, IR exposed, or IR exposed with concurrent AZD4573 (40 nM) (n=2). Genome is clustered by change in H3K27ac ChIP-seq peaks following IR exposure. Metagene profile for each cluster is on right, with profile of the dominant cluster emphasized. **e**. H3K27ac metagene profile at transcripts differentially upregulated following IR in the presence or absence of AZD4573. **f**. p-Pol II heatmap of untreated controls, IR exposed, or IR exposed with concurrent AZD4573 (40 nM) (n=3). **g**. Histogram of IR-induced transcripts LFC value in the presence or absence of AZD4573. **h**. IR-induced gene LFC value in the presence or absence of AZD4573 in SF8628 and HSJD-DIPG007 cells.

We then repeated the ATAC-seq, H3K27ac ChIP-seq, p-Pol II CUT&RUN, and RNA-seq experiments in the presence of AZD4573 co-treatment (n=2-3, 40 nM dosed 2 hours prior to irradiation) and examined these in comparison to our delineated IR-induced reorganization. Remarkably, the ATAC-defined chromatin compaction observed at both enhancers and promotors after radiation was almost completely abrogated in the presence of concurrent PTEFb inhibition (**Figure 3c**). The small subset of loci which gained accessibility following radiation were comparatively unaffected by AZD4573 treatment. Comparison of differential ATAC-seq peaks between IR- and AZD+IR-treated conditions identified 2,960 differential loci, with ontology analysis yielding similar enrichment in programs related to stress response, transcriptional processing, and cell cycle regulation (**Extended Data Table 1c**). This suggests that the adaptive chromatin compaction observed following IR not only occurs independent of a transcriptional regulatory function, but that it is instead dependent on transcriptional activity itself.

In contrast, we observed a different pattern of response in H3K27ac occupancy with AZD4573 co-treatment. At loci which lost acetylation following IR exposure, little change was seen with PTEFb inhibition. However, at promoter regions which gained H3K27ac after IR, acetylation paradoxically was markedly increased with AZD4573 co-treatment (**Figure 3d**). Enhancer elements with H3K27ac gains following IR, though small in number, showed a more modest increase in acetylation with co-treatment. Given the mechanistic link between this epigenetic modification and transcriptional output, we again examined H3K27ac occupancy and p-Pol II elongation specifically at the TSS of transcripts differentially upregulated following IR. PTEFb inhibition resulted in a significant accumulation of H3K27ac deposition immediately (<500 bp) downstream from the TSS at these loci (**Figure 3e**). Despite this, active transcription from these loci was largely abolished. p-Pol II profiles showed a complete loss of downstream egress with concurrent AZD4573 treatment, instead showing a sharp peak paused at the promoter-proximal region (**Figure 3f**). Of the 1,337 unique transcripts differentially upregulated following IR (LFC ≥1.2 qval <0.05), only 230 maintained significant upregulation in the presence of AZD4573 (**Figure 3g** and **Extended Data Figure 2a**). This created a promoter state in which inhibition of CDK9-mediated pause-release decouples H3K27ac occupancy from productive transcriptional elongation, with the observed hyperacetylation presumably reflecting ineffective epigenetic upregulation accruing behind a stalled Pol II. To test the reproducibility of this phenomenon across different model systems, we performed RNA-seq (n=3) in the same conditions in additional cell lines (SF8628 and HSJD-DIPG007) representing unique molecular backgrounds (including *H3F3A, TP53*, and *ACVR1* status). In each instance, genes with induced expression following IR exposure remained largely quiescent in the presence of AZD4573 co-treatment (**Figure 3h** and **Extended Data Figure 2b**). Together, these findings outline a model in which H3K27ac-driven recruitment of PTEFb is required for early Pol II induction following exposure to IR, and that selective inhibition of CDK9 catalytic activity within that window of time largely abrogates this adaptive response.

### PTEFb activity is required for the early induction of many canonical DNA damage response programs

To characterize the functional consequences of this imparted transcription defect, we next examined the gene expression networks enriched within the RNA-seq datasets (SU-DIPG4, HSJD-DIPG007, and SF8628). Early expression changes following a single IR exposure unsurprisingly showed enrichment in canonical DNA damage response (DDR) programs. Specific genes upregulated showed only modest overlap between models (**Extended Data Figure 3a-b** and **Extended Data Table 3**), perhaps reflecting the genomic heterogeneity between cultures (e.g. mutations to *H3F3A, HIST1H3B, TP53*, or *ACVR1*) that has been shown to contribute to non-uniformity in radiation response ^46-48^. Despite this, across the three culture systems ontology analysis of genes induced by IR exposure but abrogated in the presence of AZD4573 consistently resolved into functional programs broadly involved in transcriptional processing, DNA repair, and cell cycle regulation (**Figure 4a-b** and **Extended Data Table 4**). Alterations in transcriptional processing recapitulated our earlier observations and included enrichment in terms related to chromatin organization, transcription initiation, RNA metabolism, 3’-end processing, and RNA splicing, with specific perturbation of *CSTF1, CDK12*, and mediator subunit expression observed (**Extended Data Table 4**).

**Figure 4.**
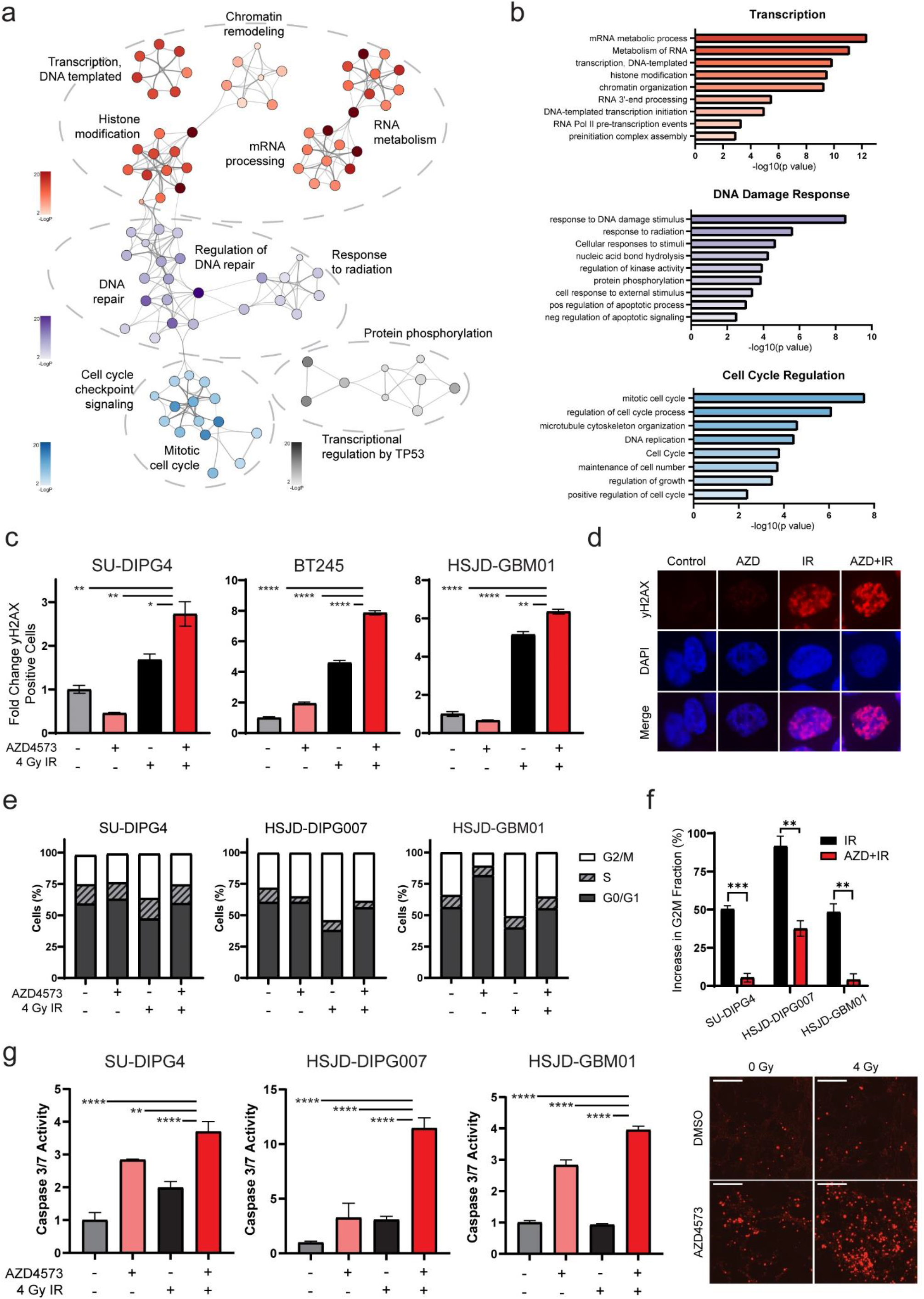
PTEFb activity is required for canonical DNA damage response programs. Gene ontology network (**a**) and terms (**b**) constructed from genes significantly induced by IR but abrogated with concurrent CDK9i across three cell lines (SU-DIPG4, HSJD-DIPG007, SF8628; n=3 each condition, LFC +/- 1.2 with p val <0.05). Each node denotes an enriched term, with color density reflecting -log(Pval). **c**. DNA damage as measured by flow cytometry for yH2AX 6 hours after IR, AZD4573, or combination. Comparison reflects two-tailed Student’s t-test (* p=<0.05, ** p=<0.01, **** p=<0.0001). **d**. Representative immunofluorescent staining for yH2AX at same timepoint and conditions as (c). **e**. Cell cycle distribution of indicated cultures after treatment with AZD4573, IR, or combination. **f**. IR-induced G2M arrest in the presence or absence of AZD4573 as measured by increase in G2M fraction from (e). Comparison reflects two-tailed Student’s t-test (** p=<0.01, *** p=<0.001). **g**. Caspase 3/7 activation measured 24 hours after treatment with AZD4573, IR, or combination. Comparison reflects two-tailed Student’s t-test (** p=<0.01, **** p=<0.0001). Representative fluorescent live-cell imaging shown on right (bar = 200 µm).

We next validated the predicted deleterious impact of PTEFb inhibition on the activation of DNA damage response programs. The phosphorylation of the histone variant H2AX can be used as an indirect marker of DNA double strand breaks, the most injurious of IR-induced DNA lesions which unrepaired can lead to genomic instability and cell death ^49^. We examined accumulation of γH2AX foci in DIPG (SU-DIPG4 and HSJD-DIPG007) and HGG (HSJD-GBM001) cultures following IR in the presence or absence of AZD4573. This revealed a reproducible increase in γH2AX-positive cells in combination-treated cultures relative to control or IR alone (**Figure 4c-d**). Arrest of the cell cycle at the G2/M checkpoint is key regulatory mechanism for ensuring genome stability after the detection of IR-induced DNA damage ^50^. Consistent with this, cell cycle distribution in DIPG and HGG cultures showed between 48-91% increase in G2/M fraction following a single IR exposure. This protective arrest was largely abolished, however, in the presence of concurrent PTEFb inhibition, with subsequent cell cycle distribution showing little to no change in comparison to untreated controls (**Figure 4e-f**). Consequently, when we examined cell death via induction of caspase 3/7 activity in these conditions, we observed a marked increase in apoptosis when cells were exposed to IR with PTEFb effectively inhibited (**Figure 4g**). Our findings suggest that much of the canonical protective cellular response to IR-induced DNA damage is contingent on an early PTEFb-mediated transcriptional engagement, and that concurrent CDK9i may cause these critical adaptive programs to collapse.

### Concurrent PTEFb inhibition exhibits cytotoxic synergy with ionizing radiation

Given the observed functional consequences to canonical pro-survival programs, we examined whether CDK9 pharmacologic inhibition could be employed concurrently with IR in order to augment its therapeutic effect. We first sought to optimize the timing of AZD4573 administration around IR exposure. Consistent with an early critical window for IR-adaptive transcriptional response, we found that maximal induction of apoptosis was achieved with treatment just prior to or concurrent with IR exposure; addition of AZD4573 after IR showed no additional pro-apoptotic effect (**Extended Data Figure 4a-b**). Using this approach, clonogenic survival of both DIPG (SU-DIPG4 and HSJD-DIPG007) and HGG (HSJD-GBM001) cultures was significantly diminished by AZD4573 in combination with IR when compared to either intervention alone (**Figure 5a**). We observed no difference in sensitivity to PTEFb inhibition based on histone or *TP53* mutation status (**Extended Data Figure 5a-c**). When cultures were seeded at increasing density and normalized to plating efficiency, a fixed low dosing of AZD4573 (2 nM) resulted in a significant radiation dose enhancement effect consistent with synergistic (as opposed to additive) interaction (**Figure 5b**).

**Figure 5.**
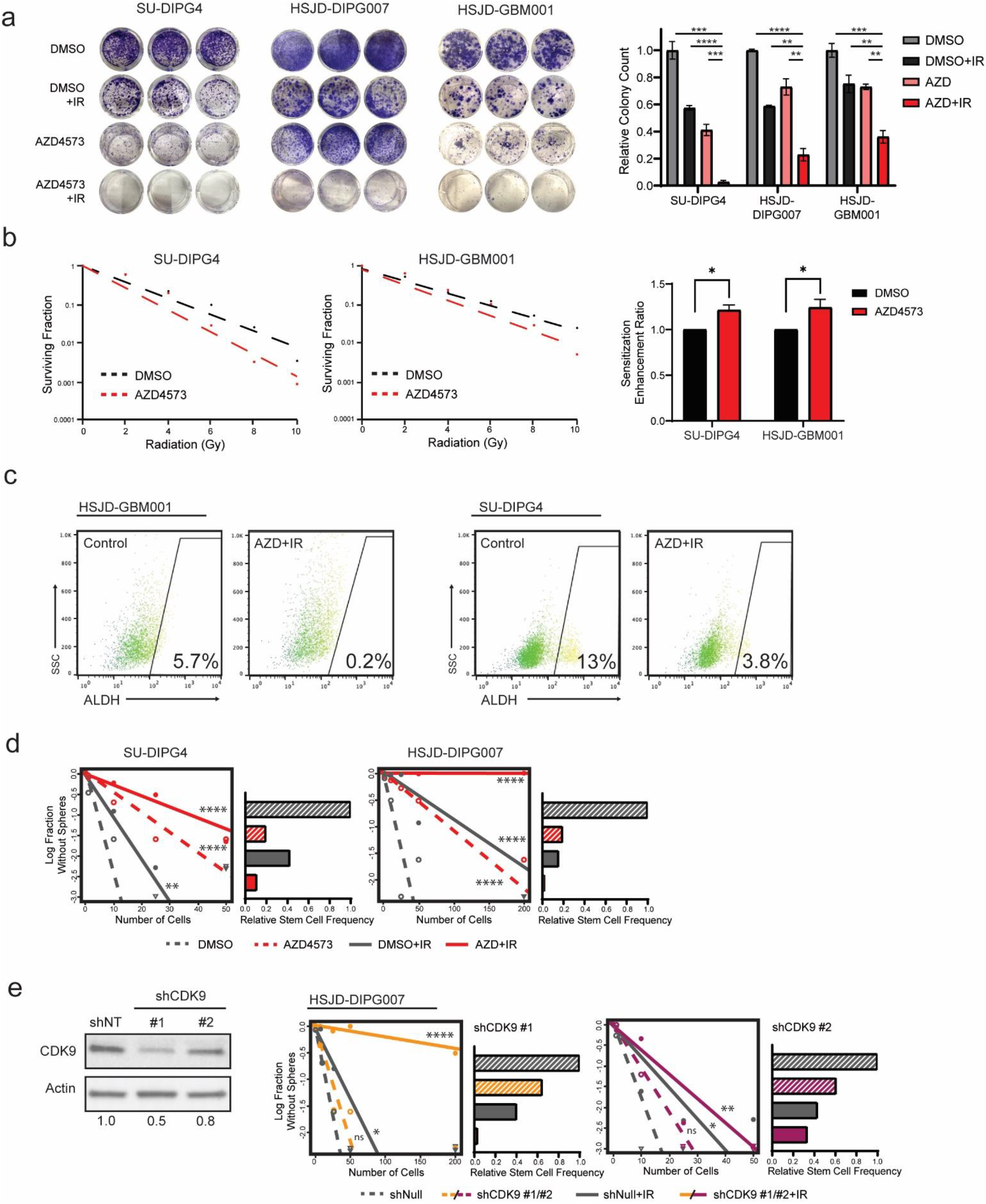
CDK9i exhibits cytotoxic synergy with IR in HGG. **a**. Colony focus assay images (left) and quantification (right) of HGG cultures treated with AZD4573, IR, or combination. Quantitative comparisons reflect two-tailed Student’s t-test (** p=<0.01, *** p=<0.001, **** p=<0.0001). **b**. Clonogenic survival (left) for HGG cultures treated with IR alone or combination AZD4573 (2 nM) + IR. Sensitization enhancement ratio (right), shown as mean + SEM, from triplicate samples normalized to plating efficiency (two-tailed Student’s t-test, * p=<0.05). **c**. Brain tumor initiating cell fraction after combinatorial AZD4573 + IR treatment as identified by ALDH expression. **d**. Neurosphere formation efficacy (left) and relative stem cell frequency (right) by extreme limiting dilution assay following treatment with AZD4573, IR, or combination. Comparisons reflect pairwise Chi-square test for stem cell frequencies (** p=<0.01, **** p=<0.0001). **e**. Western blot analysis (left) for CDK9 following shRNA transduction, relative densitometry quantification shown below. Neurosphere formation efficacy and relative stem cell frequency by extreme limiting dilution assay following shRNA transduction shown on right (pairwise Chi-square * p=<0.05, ** p=<0.01, **** p=<0.0001).

Orthogonal to measurements of gross viability or proliferation is assessment of the brain tumor initiating cell (BTIC) fraction within a tumor population. These stem-like cells are responsible for tumor initiation and have been linked to therapy resistance and cancer regrowth after treatment ^51-53^. Aldehyde dehydrogenase (ALDH) has been proposed as a BTIC marker in both pediatric and adult gliomas ^54,55^. We quantified ALDH expression after AZD4573, IR, or combination and observed a significant depletion of the ALDH+ cell fraction following combinatorial therapy (**Figure 5c** and **Extended Data Figure 6**). We next assessed potential for self-renewal directly through neurosphere extreme limiting dilution assays (ELDA). This demonstrated a reduction in stem cell frequency within all treatment arms but most significantly in combination treated cells (**Figure 5d**). To confirm that this effect was a result of AZD4573’s action against CDK9 specifically, we performed parallel experiments using shRNA knockdowns targeting CDK9. This replicated the previously observed reduction in stem cell frequency, with the degree of reduction paralleling shRNA knockdown efficiency (**Figure 5e**). Collectively, these data support the use of CDK9 inhibition as a means of augmenting the established therapeutic effect of ionizing radiation.

**Figure 6.**
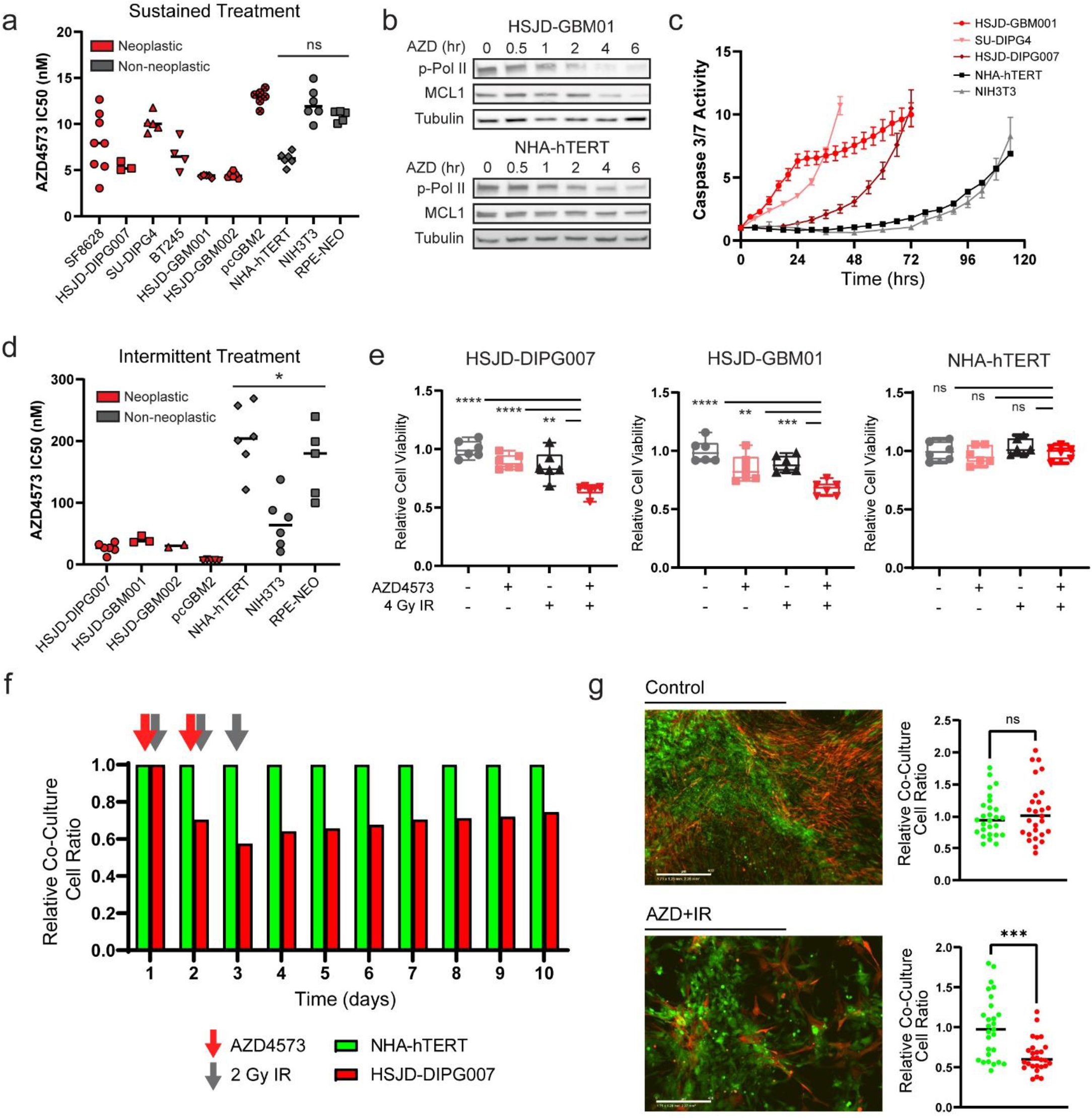
Transcriptional addiction in HGG gives rise to a therapeutic index for CDK9i relative to normal astrocytes. **a**. Half-maximal inhibitory concentration of AZD4573 after 3-day exposure in respective cell lines. Comparison reflects *p* value of two-tailed Student’s t-test of mean IC50 values from neoplastic vs non-neoplastic cultures. **b**. Western blot analysis of p-Pol II (Ser 2) and MCL1 after indicated exposure times to 50 nM AZD4573. **c**. Caspase 3/7 activity over time following fixed dose of AZD4573. Error bars indicate SEM from minimum 3 biological replicates. **d**. Half-maximal inhibitory concentrations and comparison as in (A) but measured 3 days after a single 8-hour drug exposure followed by drug washout. **e**. Cell viability measured at 3 days following 8-hour exposure to 5 nM AZD4573 +/- 4 Gy IR. Box whiskers represent min to max range of replicates. Comparison reflects two-tailed Student’s t-test (** p=<0.01, *** p=<0.001, **** p=<0.0001). **f**. Relative ratio of co-cultured DIPG cells (HSJD-DIPG007) and normal astrocytes (NHA-hTERT) following fractionated radiotherapy and intermittent AZD4573 treatment as indicated by arrows. **g**. Quantification of (f) at day 10, two-tailed Student’s t-test p=0.0001.

### Relative transcriptional addiction in HGG supports a therapeutic index for CDK9i in comparison to non-transformed cells

One could reasonably anticipate that sustained inhibition of transcriptional elongation would be uniformly deleterious across most cellular systems, raising the potential for unacceptable toxicities when adapted for clinical use. Neither of the PTEFb-member CDK9/CyclinT pair are recurringly mutated or overexpressed in pediatric HGG ^56-58^. In fact, CDK9 has been characterized as a pan-essential gene by several groups, with a loss of fitness or cell death observed following inhibition in multiple normal tissues or human cell lineages ^59-61^. Consistent with this, when we broadened our cell viability assays following 72 hours of continuous exposure to AZD4573, we observed no difference in selectivity between neoplastic glioma cultures and normal cell controls (e.g. astrocytes, fibroblasts, or epithelial cells) (**Figure 6a**).

Despite this, the pharmacologic targeting of pan-essential genes has formed the core of most successful systemic chemotherapeutic regimens, so long as proper consideration is given to strategies maximizing a therapeutic index between normal tissues and a target cell population of interest ^59^. Transcriptional addiction can be defined as an acquired reliance on the continuous activity of an oncogenic transcriptional program ^62^. This phenomenon has been identified spanning many cancer types and driver mutations, and it gives rise to specific transcriptional dependencies in cancer cells which are comparatively absent in their non-transformed counterparts ^62-67^. We examined both Pol II Ser2 phosphorylation, the primary catalytic output of PTEFb ^21^, as well as MCL1 expression, a rapid-turnover anti-apoptotic protein with established PTEFb-dependence ^45^, over time following AZD4573 treatment in both glioma and normal astrocyte cultures. At equivalent AZD4573 dosing and timepoints, we observed a more rapid abrogation of PTEFb catalytic activity and depletion of MCL1 expression in HGG cells when compared to normal astrocytes (**Figure 6b**). This correlated with apoptosis as assessed by caspase 3/7 activity on live cell imaging; while all culture systems eventually displayed comparable evidence of cell death, induction was markedly more rapid in neoplastic models in comparison to normal controls (**Figure 6c**). In light of this time-dependent differential sensitivity, we transitioned to a short-term exposure strategy for AZD4573. This was modeled *in vitro* by drug washout after 8 hours, which mimicked the rapid target engagement and established half-life from PK/PD studies *in vivo* ^45^. This strategy revealed a significant difference in sensitivity between neoplastic and non-transformed culture models (**Figure 6d**), supportive of a therapeutic index amenable to tolerable intervention.

Normal brain tissue exhibits a relative radioresistance when compared to most glial neoplasms ^68,69^, a feature exploited daily in the routine clinical treatment of CNS malignancies ^70^. Given the differential sensitivity to CDK9i we observed using an intermittent dosing strategy, we examined the comparative tolerance of normal astrocytes to this combinatorial regimen. Using a fixed dose of AZD4573 and IR, we were able to elicit a significant decrease in DIPG and GBM culture viability while sparing parallel normal astrocyte and fibroblast controls (**Figure 6e** and **Extended Data Figure 7**). To further substantiate this therapeutic index, we generated a co-culture system in which DIPG cells (HSJD-DIPG007) and astrocytes (NHA-hTERT) were transfected with GFP- or NucRed-expressing lentivirus, respectively, before plating together at equivalent density. Following fractionated radiotherapy (2 Gy x 3 doses) combined with intermittent AZD4573 administration, we observed a selective depletion of the DIPG fraction relative to normal astrocytes (**Figure 6f-g** and **Extended Data Figure 8**). Together, these data support the existence of overlapping therapeutic windows in which differential sensitivities to CDK9i and radiation could be exploited to achieve anti-tumor effect while minimizing toxicities to normal CNS tissue.

**Figure 7.**
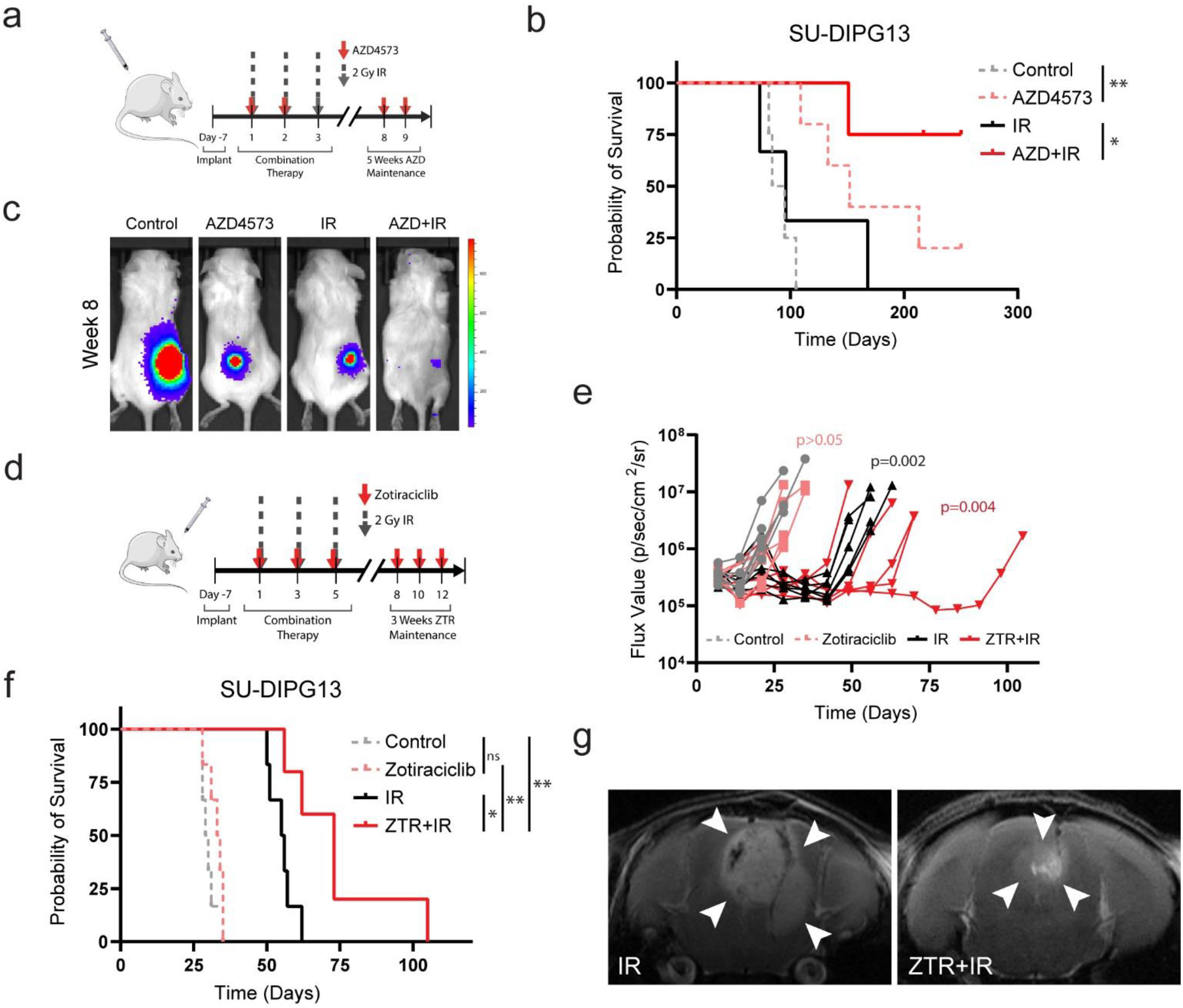
Concurrent CDK9i augments anti-tumor effect of IR to prolong survival *in vivo*. **a**. Schematic represents the treatment schedule of SU-DIPG13* xenografts with either AZD4573 (15/15 mg/kg biweekly administered intraperitoneally), radiotherapy (2 Gy x 3 fractions), or combination. **b**. Kaplan-Meier survival analysis of SU-DIPG13* flank cohorts receiving indicated treatments. **c**. Bioluminescent imaging from median mouse of each treatment cohort in (B) at completion of therapy period. **d**. Schematic representation of schedule for SU-DIPG13* xenografts treated with either zotiraciclib (ZTR, 50 mg/kg 3x weekly for two weeks followed by 35 mg/kg 3x weekly for two weeks, administered by oral gavage), radiotherapy (2 Gy x 3 fractions), or combination. **e**. Bioluminescent flux from individual animals within indicated treatment groups (Mann-Whitney test at completion of treatment, p value in insert). **f**. Kaplan-Meier survival analysis of orthotopic xenograft cohorts receiving indicated treatments. **g**. Representative T2-weighted turboRARE axial MRI sequences of IR- or ZTR+IR-treated mice. Arrowheads indicate margins of tumor.

**Figure 8.**
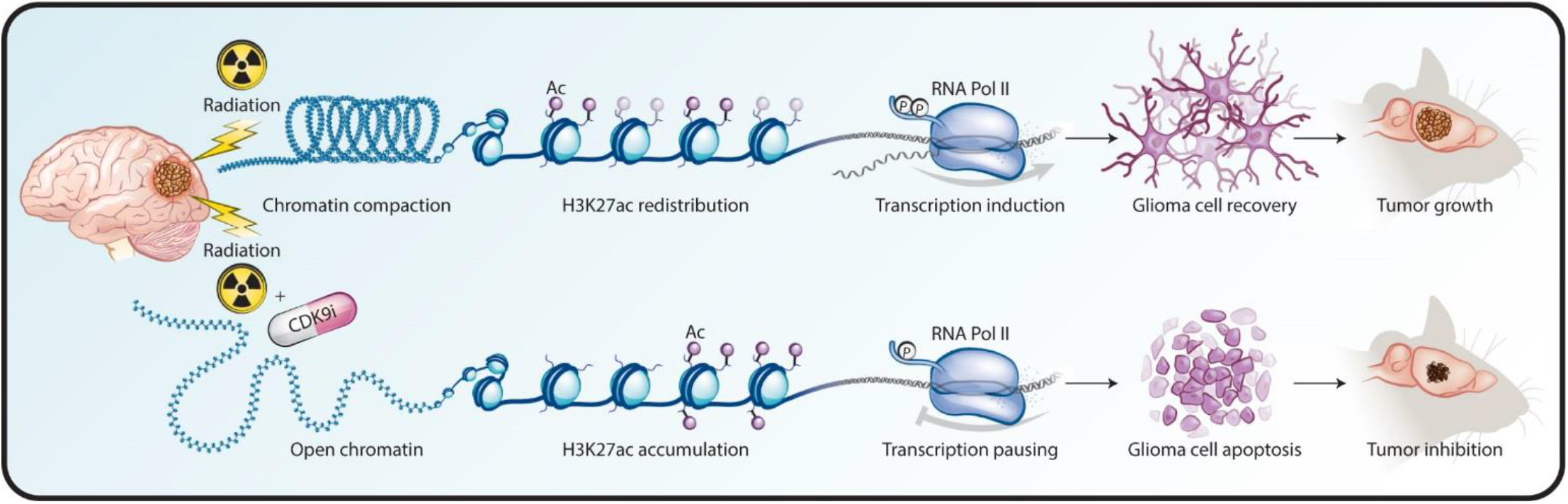
Model of IR-induced transcriptional reorganization. Rapid reorganization of active chromatin drives transcriptional induction required for DDR programs. Concurrent inhibition of PTEFb-mediated transcriptional elongation abrogates this adaptive response, augmenting the anti-tumor effect of radiotherapy.

### Effective PTEFb inhibition augments the anti-tumor effect of radiotherapy in murine models of HGG

Finally, we examined whether this approach of disrupting IR-adaptive transcriptional circuitry could be employed for therapeutic effect in animal models of pediatric high-grade glioma. We utilized SU-DIPG13*, a well-characterized, aggressive model of DIPG, engrafted in the flank of NOD scid gamma mice. Treatment with AZD4573 (15/15 mg/kg biweekly ^45^), either alone or in combination with radiotherapy (2 Gy x 3 doses), resulted in a marked delay of tumor progression and prolongation of survival, including long-term survival in a substantial portion of animals (**Figure 7a-c**). Unfortunately, AZD4573 is likely to be a substrate of both the ABCB1 (P-glycoprotein/MDR1) and ABCG2 (breast cancer resistance protein [BCRP]) cell membrane transporters ^71,72^ based on *in vitro* transporter assays, with a high efflux ratio predicted across the blood-brain barrier (BBB) (AstraZeneca, Investigator Brochure). Consistent with this, the SU-DIPG13* engrafted orthotopically demonstrated no survival benefit from treatment with AZD4573, either alone or in combination with radiotherapy (**Extended Data Figure 9a-b**).

Zotiraciclib is an orally-available CDK9 inhibitor with established CNS penetrance in several preclinical models ^73-75^ and an acceptable safety profile in phase I/II trials for adults with anaplastic astrocytoma or glioblastoma (^76^ and NCT03224104, NCT02942264). Within our culture models of pediatric HGG, zotiraciclib replicated previous CDK9-dependent effects on Pol II phosphorylation, MCL1 depletion, and radiosensitization *in vitro* (**Extended Data Figure 10**). We then tested zotiraciclib (50/35 mg/kg 3x weekly) in combination with fractionated radiotherapy (2 Gy x 3 doses) against the SU-DIPG13* model engrafted orthotopically in the pons (**Figure 7d**). This resulted in a decrease in tumor growth by bioluminescent imaging and a significant prolongation of survival in animals treated with combination therapy compared to radiotherapy alone (**Figure 7e-g**).

In total, these findings replicate our *in vitro* observations that effective inhibition of PTEFb catalytic activity augments the anti-tumor therapeutic effect of IR. While the unique anatomic and physiologic considerations for CNS tumors necessitate the selection of appropriate agents for adequate CNS delivery, our data supports the use of clinically-relevant CDK9 inhibitors in a multimodal treatment approach for these lethal cancers.

## Discussion

This integrative epigenetic and transcriptional analysis provides a framework for understanding the reorganization of transcriptionally active chromatin that underlies the early adaptive response to radiotherapy. Specifically, we show that pHGG cells rapidly compact active chromatin while redistributing H3K27ac deposition in order to modulate transcriptional output in the early hours following an IR exposure. Importantly, we identify that key enzymatic steps in this adaptive cascade are amenable to pharmacologic targeting. Inhibition of the CDK9-catalyzed phosphorylation of the Pol II CTD during the immediate peri-radiation window abrogates not only the induction of critical transcriptional DDR programs, but also significant components of the chromatin reorganization itself (**Figure 8**). This inhibition demonstrates a marked anti-tumor effect agnostic of tumor histone, TP53, or passenger mutation status, opening a therapeutic avenue that may prove resilient to the intra- and intertumoral heterogeneity that has complicated translational efforts in a molecularly diverse disease entity.

Pediatric high-grade gliomas are highly lethal malignancies, accounting for the largest proportion of cancer-associated deaths in children ^77^. Radiation remains the only uniformly accepted standard of care across HGG subtypes. As such, mechanisms to further sensitize these tumors to radiotherapy, augmenting the depth and duration of clinical response, remain attractive strategies for clinical practice (reviewed elegantly in Metselaar, et al^48^). Unlike the distributed toxicity potential with combinatorial systemic therapies, the synergistic potential of a radiosensitizing agent is largely confined to the conformal radiation field. A synergistic interaction with standard treatment also lowers practical barriers to clinical translation, as novel radiosensitizers can be more readily incorporated into existing therapy backbones for up-front phase 1 investigation.

Our present study builds on the body of data dissecting the complex reorganization of transcriptional machinery required for the rapid, coordinated induction of large gene expression programs in response to exogenous stressors. This has been best characterized within models of heat shock ^27,30,78-81^, in which Pol II is released from a paused state into gene bodies for productive elongation across heat shock responsive loci. More recent work has defined considerable specificity within this response. PTEFb may exit its 7SK-Hexim reservoir and localize to chromatin in several active complexes, including BRD4-PTEFb, AFF1-SEC, or AFF4-SEC ^41,82,83^. Using an auxin-inducible degron system, Zheng et al showed that AFF4-SEC was predominantly responsible for early induction of the heat shock stress response, while other active PTEFb-containing complexes were largely dispensable ^25^. PTEFb-dependent transcriptional alteration has now also been defined in stress responses to other genotoxic stimuli including chemical carcinogens, albeit with less clearly delineated complex specificity ^26^. Our approach here using short hairpin RNAs or small molecules targeting the CDK9 catalytic component of PTEFb accepts a tradeoff of molecular specificity for translational potential. Our data lacks the granularity to convincingly define which PTEFb-containing complex is primarily responsible for the observed IR-induced response, limiting extrapolation to the expected benefit from BET inhibitors ^84^ or SEC-specific compounds ^85,86^. However by targeting the common CDK9-mediated catalytic reaction directly, this strategy bypasses the redundancy observed to some degree with the various PTEFb-containing complexes ^25^, ensuring the desired therapeutic endpoint agnostic of a distinct carrier complex.

CDK9 has been recognized as a promising target for cancer therapy for more than a decade, prompting the formulation of numerous inhibitory compounds now in various stages of preclinical and clinical development ^87,88^. These have ranged from the first multi-kinase inhibitors such as flavopiridol and dinaciclib to newer, highly-selective agents like AZD4573 ^45^ or intriguing peptidomimetics targeting SEC directly ^85^. Early phase I and II clinical trials with flavopiridol and dinaciclib were limited by high rates of adverse events with only modest disease response ^89-94^, frequently attributed to dose-limiting off-target effects by these relatively non-specific compounds ^87^. Phase I/II trials with more selective agents are now ongoing (e.g. NCT03263637, NCT04630756, NCT03224104, NCT02942264). Our data here suggests that not only selectivity but timing and duration of dose exposure is critical in achieving a tolerable therapeutic index, supporting similar observations reported in the preclinical development of AZD4573 for hematologic malignancies ^45^. It likewise echoes what has now been borne out in early clinical trials, in which a phase I dose escalation study of zotiraciclib encountered significant dose limiting toxicities (DLTs) with continuous daily dosing while an intermittent dosing schedule was able to reach the maximum defined dose level without DLTs reported ^95^. Furthermore, as many CDK9 inhibitors are under active clinical development primarily for hematologic malignancies or extracranial sarcomas, few compounds have robust data available on effective penetration across the blood brain barrier. Our data here supports the broader testing of this class of compounds against CNS malignancies.

In conclusion, this study identifies PTEFb as serving a pivotal role in mediating the transcriptional reorganization required to support an early DNA damage response to radiotherapy. The centrality of this function creates a therapeutic window surrounding IR exposure, in which effective PTEFb inhibition causes many core adaptive programs to collapse. Judicious dosing strategies can safely exploit this dependency in models of high-grade gliomas, opening a new avenue of therapy for these recalcitrant malignancies.

## Methods

### Cell lines and culture technique

HGG and normal cells were maintained as previously described ^33,96^. Briefly, SU-DIPG4 and SU-DIPG13* cells were cultured in tumor stem media (TSM) consisting of Neurobasal(-A) (Invitrogen), human-basic FGF (20 ng/mL; Shenandoah Biotech), human-EGF (20 ng/mL; Shenandoah Biotech), human PDGF-AB (20 ng/mL; Shenandoah Biotech), B27(-A) (Invitrogen), and heparin (10 ng/mL). HSJD-DIPG007 cells were maintained in TSM media as above with 10% fetal bovine serum (FBS)(Atlanta Biologicals). BT245 cells were grown in NeuroCult NS-A media (Stemcell Technologies) supplemented with penicillin-streptomycin (1:100, PenStrep)(Gibco/ThermoFisher), heparin (2 μg/mL), human EGF (20 ng/mL), and human FGFb (10 ng/mL). SF8628 cells were maintained in Dulbecco’s modified eagle medium (DMEM)(Gibco/ThermoFisher) and supplemented with 10% FBS, MEM non-essential amino acids (Gibco/ThermoFisher) and antibiotic/antimicotic (Gibco/ThermoFisher). Normal human astrocytes were cultured in DMEM and supplemented with 10% FBS, PenStrep, L-glutamine (Gibco/ThermoFisher), and sodium pyruvate (Gibco/ThermoFisher). NIH3T3 cells were cultured in DMEM and supplemented with 10% FBS and PenStrep. Cells were grown in either adherent monolayer conditions (Falcon/Corning) or as tumor neurospheres (ultra-low attachment flasks, Corning) as indicated. All cell lines were validated by DNA fingerprinting through the University of Colorado Molecular Biology Service Center utilizing the STR DNA Profiling PowerPlex-16 HS Kit (DC2101, Promega)(**Extended Data Figure 11**).

### Lentiviral production

HEK293FT cells were used to produce viral particles by packaging PSPAX2 and PMD2.G vectors with shCDK9 plasmids purchased from the University of Colorado Functional Genomics Core Facility (TRCN0000000494 and TRCN0000199780).

### Sensitivity enhancement ratios

Cells were seeded in triplicate 1,000 to 10,000 cells in 6 well plates. They were treated with zero to 10 Gy at 2 Gy increments in the presence or absence of 3nM AZD4573. After 14 days, media was aspirated, cells were washed with 1x PBS, and stained with crystal violet for 15 minutes before being manually counted.

### Cell viability assays

Cells were treated with AZD4573 (AstraZeneca), TG02 (MedChemExpress), or ionizing radiation as indicated before adding MTS reagent (Promega) and measuring absorbance at 492 nM on a BioTek Synergy H1 microplate reader (BioTek Instruments, VT).

### Caspase 3/7 assays

Indicated cells were seeded at a density of 4000 cells per well on 96 well plates. Radiation was administered at 4 Gy, immediately followed by the addition of caspase 3/7 dye (Sartorius) at 0.5uM. Cells were monitored over time using IncuCyte S3 Live Cell Analysis System, with red or green reporter count normalized against time zero values.

### Aldehyde dehydrogenase assay

ALDH activity of was measured using Aldefluor kit (Stem Cell Technologies) according to the manufacturer’s instruction. Cells were stained with propidium iodide and then analyzed on the Guava easyCyte HT flow cytometer (Luminex).

### Extreme limiting dilution assay

Limiting dilution assays were performed as we ^33^ and others ^97^ have previously described. Briefly, cells were seeded on a 96-well ultra-low-attachment round-bottom tissue culture plate in serum-free media at increasing numbers from 1 cell/well to 100 cells/well. Cells were allowed to grow for 14 days, and the number of wells containing neurospheres was counted manually under light microscopy. Published ELDA software (http://bioinf.wehi.edu.au/software/elda/) was used to calculate comparative self-renewal potential of cells.

### H2AX flow cytometry

Cells were seeded and radiated in the presence or absence of AZD4753. At least 1,000,000 cells were collected per condition and washed with 2% FBS in 1x PBS wash buffer. The DMSO control was split into two samples to be used as a negative control. Cells were spun and resuspended in 100uL of BD CytoFix Fixation buffer (554655) for 10 minutes at room temperature. Samples were spun, washed, and incubated with -20-degree BD Phosflow permeation buffer (558050) for 5 minutes at room temperature. After incubation, cells were spun and washed again and resuspended in Alexa Fluor 488.

### Click-iT nascent RNA imaging

Cells were plated at 50% confluency in a 96-well microplate. The following day, cells were irradiated at 4 Gy and proceeded to the reaction immediately after the radiation, 1 hr, 2 hrs, 4 hrs, and 8 hrs after radiation, or treated with 0, 10, 20, 30, or 40 nM of AZD4573 for 1 hour. Reactions were performed accordingly to the manufacturer instructions (Click-iT RNA HCS Assay, C10327, Invitrogen). Briefly, cells were treated with 1mM of EU and incubated under normal cell conditions for 1 hour. After EU incubation, media was removed and 50 ul of 3.7% formaldehyde in PBS in each well was added and incubated for 15 min at room temperature. The fixative was then removed, and cells were washed once with PBS. 50 ul of 0.5% Triton X-100 in PBS was added and incubated for 15 minutes at room temperature. Cells were washed once with PBS, and 50 ul of Click-iT cocktail was added and incubated for 30 minutes at room temperature, protected from light. After the incubation, Click-iT cocktail was removed and cells were washed once with 50 ul of Click-iT reaction rinse buffer. Cells were washed with PBS and then incubated with 50 ul of HCS NuclearMask Blue stain solution, diluted 1:1000 in PBS, and incubated for 15 minutes at room temperature, protected from light. HCS NuclearMask Blue stain solution was removed and cells were washed with PBS twice. Fluorescence was then quantified using the Incucyte S3 (Sartorius).

### Cell cycle analysis

Cells were seeded 500,000 per 100mm dish and treated at 70 - 80% confluence with DMSO, 8nM AZD5473, 4Gy radiation, or combination. Cells were collected at 24 hours, fixed with 1-3mL ice cold 70% EtOH and stored at -20 degrees overnight. The pellet was washed in a wash buffer of 2% FBS in 1x PBS, spun, and resuspended in 100uL of fluorescent nuclear dye DRAQ5, diluted 1:1000 in wash buffer. The suspensions were incubated for 30 minutes at 37-degrees in the dark. After incubation, cells were run on an Amnis FlowSight cytometer equipped with a 488 nm laser.

### Western blotting

Whole-cell protein lysates were harvested in lysis buffer (RIPA buffer supplemented with protease inhibitor (Roche), sodium vanadate and sodium molybdate) from cells in indicated conditions. Protein was separated on 4-20% PROTEAN TGX Gels and blotted using a wet transfer system (Biorad) before probing for CDK9 (Abcam [EPR3119Y] (ab76320)), phospho-Rbp1 CTD (Ser2) (Cell Signaling (E1Z3G) #13499), phospho-Rbp1 CTD (Ser5) (Cell Signaling (D9N5I) #13523S), or α-Tubulin (Cell Signaling (DM1A) #3873S).

### Immunofluorescence

Cells were plated in 8-well chamber and 24 hours later were submitted to the following treatments: DMSO, 8nM AZD, IR, or combination. IR was given 2 hr after drug treatment. Reaction for γH2AX (BD Pharmigen; Alexa Fluor 488 Mouse anti-H2AX (Ps139); 560445) was performed 6hs after radiation; reaction for CDK9 (Cell Signaling, (C12F7) #2316S) was performed after 4 hrs. Cells were washed in PBS, fixed with 4% paraformaldehyde for 20 min, and washed 3 times with PBS for 5 min each. Cell permeabilization was performed using Triton X-100 0.3% for 5 min and unspecific reaction was blocked incubating cells with 3% BSA for 1h at room temperature. Cells were incubated with a primary antibody diluted 1:100 in 3% BSA and incubated overnight at 4 C. For anti-H2AX reactions, cells were washed with PBS 3 times and slides were mounted with coverslips using the ProLong Gold antifade reagent with DAPI (Invitrogen; P36935). For anti-CDK9 reactions, cells were washed 3 times with PBS and secondary antibodies were applied diluted 1:1000 in PBS and incubated for 1 hr at room temperature, protected from light. Cells were washed with PBS 3 times, slides were mounted with coverslips using the ProLong Gold antifade reagent with DAPI (Invitrogen; P36935), and images were taken using the a confocal microscope (Keyence).

### Transcriptome sequencing (RNA-seq)

Ribonucleic acid was isolated from cells in indicated experimental conditions using a Qiagen miRNAeasy kit (Valencia, CA). Illumina Novaseq 6000 libraries were prepared and sequenced by the Genomics and Microarray Core Facility at the University of Colorado Anschutz Medical Campus. Resulting sequences were filtered and trimmed, removing low-quality bases (Phred score <15), and analyzed using a custom computational pipeline consisting of gSNAP for mapping to the human genome (hg38), expression (FPKM) derived by Cufflinks, and differential expression analyzed with ANOVA in R. Output files contained read-depth data and FPKM expression levels for each sample, and when gene expression levels were compared between groups of samples, the ratio of expression in log2 format and a P value for each gene was recorded. Subsequent gene ontology analysis was performed using the Metascape platform ^98^.

### Assay for transposase accessibility to chromatin (ATAC-seq)

Cells were plated in a 10 cm-plate and the following day were treated with DMSO, 40 nM AZD4573, 6 Gy IR, or combination. IR was administered 2 hours after drug treatment. 4 hours after IR, cells were scraped and counted. 100,000 cells were spun down at 500 xg for 5 min at 4°C, pellet was washed once with 500 ul of cold 1x PBS, and spun down at 500 xg, 5 min at 4°C. Pellet was resuspended in 450 ul of cold hypotonic buffer (10mM Tris-HCl, pH 7.4, 10mM NaCl, 3mM MgCl_2_) and immediately added 50 ul of 1% IGEPAL CA-630 (0.1% final) and inverted to mix. Cells were incubated on ice for 15 min, spun down at 500 xg for 10 min at 4° C. Pellet was set on ice, and the transposition reaction was performed following the manufacturer instructions (Cell Biologics, Cat No. CB6936). Pelleted cells were resuspended in 50 ul of transposition reaction mix (25 ul 2X Reaction Buffer, 2.5ul Transposome, 22.5ul Nuclease free water) and incubated at 37°C for 1 hour. DNA was purified using Qiagen MinElute kit (Qiagen, #28004) and eluted in 15 ul of Elution Buffer, followed by the Library Generation to amplify transposed DNA fragments. For this reaction the following reagents were mixed in a PCR tube, 10ul of Transposed DNA, 10 ul of Nuclease free water, 2.5 ul of Ad1.noMX (Oligo 1), 2.5 ul Ad2._Barcode (Oligo 2), and 25 ul of High Fidelity 2x PCR Master Mix. This mixture was run on a thermocycler using the following cycle: 98°C 30 sec, (98°C 10 sec, 63°C 30 sec, 72°C 1 min) x 10 times and held at 4°C. For double-sided bead purification (to remove primer dimers and large >1,000 bp fragments), each PCR sample was transferred to a 1.5 ml tube and 0.5X volume (25 ul) of AMPure XP beads was added, mixed by pipetting 10 times, and incubated at room temperature for 10min. Tubes were placed in a magnetic rack for 5 min, and the supernatant was transferred to a new tube. 1.3X of the original volume (65 ul) of AMPure XP beads was added, mixed by pipetting 10x, and incubated at room temperature for 10 min. Tubes were placed in a magnetic rack for 5 min, supernatant was discarded, and beads were washed with 200 ul 80% EtOH. EtOH was removed and tubes were left on magnetic rack with cap open for 10 min or until the pellet was totally dry. Beads were resuspended in 20ul of nuclease-free water and mixed thoroughly, tubes were placed in the magnetic rack for 5 min, and the supernatant was transferred to a new tube. Libraries were paired-end sequenced on NovaSEQ 6000 platform.

### Chromatin immunoprecipitation (ChIP-seq)

Cells were crosslinked with 1% formaldehyde added to the growth media for 10 min, followed by quenching with 0.125M glycine for 5 min. Cells were washed twice with cold PBS, collected in ice-cold PBS by scraping, pelleted, and resuspended in cell lysis buffer (5 mM PIPES, pH 8.0; 85mM KCl; 0.5% NP-40). Following incubation on ice for 10 min, a nuclear-enriched fraction was collected by centrifugation for 5min at 2500 xg at 4°C. Pellet was resuspended in ChIP lysis buffer (50mM Tris-HCl, pH 8.0; 10mM EDTA; 1%SDS) containing COmplete EDTA free protease inhibitors (Sigma), and quantified with BCA. 2 ug/ul of lysates were sonicated with a Bioruptor Plus (Diagenode) for 20 cycles (30 seconds ON and 30 seconds OFF). The size of the sonicated DNA fragments was checked on an 1X TAE 1% agarose gel electrophoresis and ranged between 250 to 500 bp. After removing the debris via centrifugation, chromatin extracts were collected.

For immunoprecipitation, chromatin extracts were diluted with Chip Dilution Buffer (16.7mM Tris-HCl, pH8.1, 1.2Mm EDTA, 167mM NaCl, 0.01%SDS and 1.1% Triton x100) and incubated with primary antibodies (anti-H3K27Ac, Active Motif #39133) overnight at 4º C. After incubation with the primary antibody, 20 uL of pre-washed magnetic beads (Magna ChIP Protein A+G Magnetic Beads, Millipore Sigma) were added to each sample for 2 hours at 4º C. Using a magnetic rack, the immunoprecipitates were washed successively with 1 ml of low salt buffer (20 mM Tris-HCl [pH 8.0], 150 mM NaCl, 0.1% SDS, 1% triton X-100, 2 mM EDTA), high salt buffer (20 mM Tris-HCl [pH 8.0], 500 mM NaCl, 0.1% SDS, 1% triton X-100, 2 mM EDTA), LiCl washing buffer (10 mM Tris-HCl [pH 8.0], 250 mM LiCl, 1.0% NP40, 1.0% deoxycholate, 1 mM EDTA) and twice with TE buffer. The DNA-protein complexes were eluted with 300 µl of IP elution buffer (1% SDS, 0.1 M NaHCO3). The cross-links were reversed by adding NaCl (a final concentration 0.2 M) into the eluents and incubating them at 65º C overnight. The DNA was recovered by proteinase K and RNase A digestion, followed by phenol/chloroform extraction and ethanol precipitation. Pellets were resuspended in 25ul of RNase and DNase free water, and ChIP-DNA was quantified using the Qubit dsDNA High Sensitivity Assay kit (Thermo Fisher Scientific).

### Cleavage under target and release using nuclease (CUT&RUN)

Beads were prepared using 10 ul/sample of CUTANA Concavalin A Conjugated Paramagnetic Beads (EpiCypher, SKU:21-1401). Beads were transferred to a 1.5 ml tube and placed in a magnetic separation rack, supernatant was removed, and beads were washed 2 times with 100 ul/sample of cold Bead Activation Buffer (20mM HEPES pH7.9, 10mM KCl 1mM CaCl_2_, 1mM MnCl2). Beads were then resuspended in 10 ul/sample of cold Bead Activation Buffer, and aliquot 10 ul/sample of activated bead slurry into 8-strip tube and kept on ice.

Cells were collected using 0.05% trypsin, washed with 100 ul/sample of Wash Buffer (20mM HEPES pH7.5, 150mM NaCl, 0.5mM Spermidine, 1x Roche cOmplete EDTA-free Protease Inhibitor (1187358001)) at room temperature, and centrifuged at 600g for 3min, for a total of two washes. Cells were resuspended in 100 ul/sample RT Wash Buffer, and then an aliquot of 100 ul washed cells was added to each 8-strip tube containing 10 ul of activated beads and mixed by pipetting. Cells were incubated with bead slurry for 10 min at RT. Tube strip was placed on a magnetic separation rack until slurry clears and the supernatant was removed. 50 ul of cold Antibody Buffer was added to each sample and mixed by pipetting; 1ul of antibody IgG (Rabbit IgG Negative Control, EpiCypher, 13-0042K), Pol II (Rpb1 CTD (4H8), Cell Signaling, #2629S), and p-Pol II (pRpb1 CTD (S2), Cell Signaling, #13499S), was added to each sample, mixed and incubated on a nutator overnight at 4°C. Tube was placed on a magnet until slurry clears, and the supernatant was removed. While beads were on magnet, 250 ul of cold Digitonin Buffer was added directly onto beads of each sample and then pipetted to remove supernatant, for a total of 2 washes. 50 ul of cold Digitonin Buffer was added to each sample and mixed. 2.5ul of CUTANA pAG-MNase (20Xstock) was added to each sample, mixed, and incubated for 10min at RT. Tube was placed on a magnet and supernatant was removed. Beads were washed 2 times with cold Digitonin Buffer. Supernatant was removed, and 50 ul of cold Digitonin Buffer was added to each sample and mixed. Tubes were placed on ice, and 1 ul of 100mM CaCl_2_ was added to each sample and mixed, and then incubated on nutator 2 hrs at 4°C. 33 ul/sample of Stop buffer containing 0.5 ng/sample of E.coli spike-in DNA, was added to the samples, mixed by pipetting, and incubated for 10 min at 37°C in a thermocycler. Tubes were placed on a magnet stand until slurry cleared, and the supernatant was transferred to a 1.5ml tube. Finally, the DNA was purified using CUTANA DNA Purification Kit (EpiCypher, 14-0050) according to the manufacturer instructions, and 1 ul of DNA was used for quantification by Qubit.

### Library preparation

The NEBNext Ultra DNA Library Prep Kit for Illumina (NEB #E7645S) with Dual Index Primers (NEB #E7600) were used for library preparation with ChIP-seq.

#### NEBNext end prep

1x TE was added to the DNA to bring final volume to 50 ul and mixed with 3 ul of NEBNext Ultra II End Prep Enzyme Mix and 7 ul of NEBNext Ultra II End Prep Reaction Buffer. This was placed in thermocycler with heated lid set to 75°C and run 30 minutes at 20°C, 60 minutes at 50°C, and held at 4°C.

#### Adaptor ligation

The adaptor ligation reaction was performed with 60 ul of Prep Reaction Mixture, 2.5 ul of NEBNext Adaptor for Illumina diluted following the manufacture recommendations, 30 ul of NEBNext Ultra Ligation Master Mix, and 1 ul of NEBNext Ligation Enhancer. This was incubated at 20°C for 15 min with heated lid off. 3 ul of USER Enzyme was added to the reaction and incubated at 37 °C for 15 min with heated led set to 47°C.

#### Cleanup of adaptor-ligated DNA without size selection

1.1x of AMPure XP Beads were added to the Adaptor Ligation reaction, and samples were incubated for 5 min at room temperature. Tubes were placed on a magnetic stand to separate beads from supernatant, and after 5 min, supernatant was removed and beads were washed 2 times with 200 ul of 80% freshy prepared ethanol. Beads were air dried for up 5 min while tubes were on magnet with lid open. DNA was eluted from the beads by adding 17ul of 0.1XTE buffer.

#### PCR enrichment of adaptor-ligated DNA

To 15 ul of the Adaptor Ligated DNA fragments was added 25 ul of NEBNext Ultra Q5 Master Mix, 5 ul of Index primer i7, and 5 ul Universal PCR Primer i5. PCR amplification was performed following the cycling conditions: 98°C for 45s, 98°C 15s and 60°C 10s for 16 cycles, 72°C for 1min, and hold at 4°C. Clean up of the PCR reaction was performed using AMPure XP Beads.

### Sequencing Analysis

The quality of the fastq files was accessed using FastQC ^99^ and MultiQC ^100^. Illumina adapters and low-quality reads were filtered out using BBDuk (http://jgi.doe.gov/data-and-tools/bb-tools). Bowtie2 (v.2.3.4.3) ^101^ was used to align the sequencing reads to the hg38 reference human genome. Samtools (v.1.11) ^102^ was used to select the mapped reads (samtools view -b - q 30) and sort the bam files. PCR duplicates were removed using Picard MarkDuplicates tool (http://broadinstitute.github.io/picard/). The normalization ratio for each sample was calculated by dividing the number of uniquely mapped human reads of the sample with the lowest number of reads by the number of uniquely mapped human reads of each sample. These normalization ratios were used to randomly sub-sample reads to obtain the same number of reads for each sample using using samtools view -s. Bedtools genomecov was used to create bedgraph files from the bam files ^103^. Bigwig files were created using deepTools bamCoverage ^104^ and visualized using IGV ^105^. Peaks were called using MACS2 (v2.1.2) ^106^ using ENCODE recommendations. IDR was used to identify the reproducible peaks between the replicates ^107^. Further processing of the peak data was performed in R, using in particular the following tools: valR ^108^, and DiffBind ^109^. Average profiles and heatmaps were generated using ngs.plot ^110^.

### Orthotopic and flank xenograft models

Animal models were generated as we have previously described ^33^. Briefly, mice were anesthetized and immobilized on a Kopf Model 940 Stereotaxic Frame with a Model 923 mouse gas anesthesia head holder and Kopf 940 Digital Display. To target the pons, a mm diameter burr was drilled in the cranium using a Dremel drill outfitted with a dental drill bit at 1.000 mm to the right and 0.800 mm posterior to lambda, with cell suspension injected 5.000 mm ventral to the surface of the skull. A suspension of 100,000 cells/2 µl/injection was then slowly injected using an UltraMicroPump III and a Micro4 Controller (World Precision Instruments). Post-surgical pain was controlled with SQ carprofen. For flank model generation, 5x10^7^ cells/200 ul serum-free media are manually injected subcutaneously into mouse flank. University of Colorado Institutional Animal Care and Use Committee (IACUC) approval was obtained and maintained throughout the conduct of the study.

### Animal treatment

Mice were randomized sequentially at 6-8 days following injection. AZD4573 was prepared to 10% DMSO stock and diluted in 40% sterile water, 39% polyethylene glycol 400, and 1% Tween 80. Mice were treated with 15/15 mg/kg IP biweekly ^45^, either alone or in combination with radiotherapy (2 Gy x 3 doses) for total of 6 cycles or until reaching protocol endpoint. Zotiraciclib (TG02) was prepared to 10% DMSO stock and diluted in 0.5% methylcellulose and 1% Tween 80 in sterile water. Mice were treated with 50 mg/kg three times weekly for two weeks followed by 35 mg/kg three times weekly for two weeks, either alone or in combination with radiotherapy. Conformal radiation was administered as previously described ^111^. Briefly, each mouse was anesthetized and positioned in the prone orientation aligned to the isocenter in two orthogonal planes by fluoroscopy. Each side of the mouse brain received half of the dose, which was delivered in opposing, lateral beams. Dosimetric calculation was done using a Monte-Carlo simulation in SmART-ATP (SmART Scientific Solutions B.V.) for the fourth ventricle, mid brain, and pons receiving the prescribed dose. Treatment was administered using a XRAD SmART irradiator (Precision X-Ray) using a 225 kV photon beam with 0.3 mm Cu filtration through a circular 10-mm diameter collimator.

### Animal imaging

Non-invasive magnetic resonance imaging was performed through the University of Colorado Anschutz Medical Campus Animal Imaging Shared Resource as previously described^33^. Scans were performed on an ultra-high field Bruker 9.4 Tesla BioSpec MR scanner (Bruker Medical, Billerica, MA) equipped with a mouse head-array RF cryo-coil. Non-gadolinium multi-sequential MRI protocol was applied to acquire (i) high-resolution 3D T2-weighted turboRARE; (ii) sagittal FLAIR; and (iii) axial fast spin echo DWI. All MRI acquisitions and image analysis were performed using Bruker ParaVision 360NEO software. All MRI acquisitions and image analyses were performed by a radiologist blinded to the treatment assignment of the mice.

### Quantification and statistical analysis

Unless indicated otherwise in the figure legend, all in vitro data are presented as mean ± SEM. In vitro assays were performed with a minimum of three independent samples, and key experiments were successfully replicated with independent samples on separate days. For quantitative comparisons, significance was defined as n.s. p>0.05, * p=<0.05, ** p=<0.01, *** p=<0.001, **** p=<0.0001. All statistical analysis was performed on GraphPad Prism 9.0 software (GraphPad, La Jolla, CA). Kaplan-Meier survival curve comparisons were performed by log-rank (Mantel-Cox) test using GraphPad Prism 9.0 software.

### Data availability

The accession number for the raw and processed data reported in this paper is GEO: GSE211149.

## Acknowledgements

We would like to thank the following individuals for generously providing the cell lines used for this work: Michelle Monje (Stanford University) for SU-DIPG4 and SU-DIPG13, Angel Montero Carcaboso (Sant Joan de Déu) for HSJD-DIPG007, HSJD-GBM001, and HSJD-GBM002, Siddhartha Mitra (Stanford University, University of Colorado) for SU-pcGBM2, and Nalin Gupta (University of California, San Francisco) for SF8268 and SF7761. We likewise thank the University of Colorado Functional Genomics Facility, Genomics and Microarray Shared Resource, Research Histology Shared Resource, and Animal Imaging Shared Resource for their contributions to this work. Scientific illustrations by Katie Vicari.

This work was generously supported by grants through the St. Baldrick’s Foundation (Fellowship Award 606095, N.A.D. and R.V.), the National Institute of Child Health and Human Development (5K12HD068372-09, N.A.D.), the National Institute of Neurological Disorders and Stroke (1K08NS121592-01A1, N.A.D.), the Morgan Adams Foundation (N.A.D., S.V., and R.V.), the Andrew McDonough B+ Foundation (N.A.D.), the Uncle Kory Foundation (N.A.D.), and the Luke Morin Family (N.A.D., S.V., and R.V.). The University of Colorado Research Histology Shared Resource is supported by a Cancer Center Support Grant (P30 CA046934). The University of Colorado Animal Imaging Shared Resource is supported by the NCI P30 CA046934 and NIH S10 OD023485 grants (N.J.S). S.D.K. is funded National Institute of Dental and Craniofacial Research to SDK (1R01DE028282-01, 1R01DE028529-01) and by 1P50CA261605-01. AZD4573 and TG02 were provided at no cost by AstraZeneca and Cothera Bioscience, respectively, under preclinical MTA.

## Author Contributions

L.M.S. performed the ChIP-seq, CUT&RUN, and ATAC-seq experiments. E.D. performed the ChIP-/ATAC-seq and CUT&RUN alignment and analysis. F.M.W. performed the RNA-seq experiments. B.S. and D.W. performed RNA-seq alignment and analysis. L.M.S. and F.M.W. performed flow cytometry and *in vitro* phenotypic assays. A.P. and F.M.W performed the stereotactic xenograft injections; F.M.W. and I.B. conducted the subsequent *in vivo* studies. S.D.K. designed and administered the *in vivo* radiotherapy plan. N.J.S. performed the xenograft MR imaging and was responsible for its analysis. N.K.F. procured tissue specimens and maintained IRB compliance. F.M.W., L.M.S., and N.A.D. prepared the figures and wrote the manuscript. S.V., N.K.F., R.D., R.V., and N.A.D. conceived the project, supervised all aspects of the work, and edited the manuscript.

## REFERENCES

1 Vignard, J., Mirey, G. & Salles, B. Ionizing-radiation induced DNA double-strand breaks: a direct and indirect lighting up. Radiother Oncol 108, 362–369 (2013). https://doi.org:10.1016/j.radonc.2013.06.013

2 Jackson, S. P. & Bartek, J. The DNA-damage response in human biology and disease. Nature 461, 1071–1078 (2009). https://doi.org:10.1038/nature08467

3 Sulli, G., Di Micco, R. & d’Adda di Fagagna, F. Crosstalk between chromatin state and DNA damage response in cellular senescence and cancer. Nature reviews. Cancer 12, 709–720 (2012). https://doi.org:10.1038/nrc3344

4 Pedersen, H., Schmiegelow, K. & Hamerlik, P. Radio-Resistance and DNA Repair in Pediatric Diffuse Midline Gliomas. Cancers (Basel) 12 (2020). https://doi.org:10.3390/cancers12102813

5 Pilié, P. G., Tang, C., Mills, G. B. & Yap, T. A. State-of-the-art strategies for targeting the DNA damage response in cancer. Nat Rev Clin Oncol 16, 81–104 (2019). https://doi.org:10.1038/s41571-018-0114-z

6 Lord, C. J. & Ashworth, A. The DNA damage response and cancer therapy. Nature 481, 287–294 (2012). https://doi.org:10.1038/nature10760

7 Goldstein, M. & Kastan, M. B. The DNA damage response: implications for tumor responses to radiation and chemotherapy. Annu Rev Med 66, 129–143 (2015). https://doi.org:10.1146/annurev-med-081313-121208

8 Rashi-Elkeles, S. et al. Parallel profiling of the transcriptome, cistrome, and epigenome in the cellular response to ionizing radiation. Sci Signal 7, rs3 (2014). https://doi.org:10.1126/scisignal.2005032

9 Liakos, A., Konstantopoulos, D., Lavigne, M. D. & Fousteri, M. Continuous transcription initiation guarantees robust repair of all transcribed genes and regulatory regions. Nat Commun 11, 916 (2020). https://doi.org:10.1038/s41467-020-14566-9

10 Venkata Narayanan, I. et al. Transcriptional and post-transcriptional regulation of the ionizing radiation response by ATM and p53. Sci Rep 7, 43598 (2017). https://doi.org:10.1038/srep43598

11 Klemm, S. L., Shipony, Z. & Greenleaf, W. J. Chromatin accessibility and the regulatory epigenome. Nat Rev Genet 20, 207–220 (2019). https://doi.org:10.1038/s41576-018-0089-8

12 Li, B., Carey, M. & Workman, J. L. The role of chromatin during transcription. Cell 128, 707–719 (2007). https://doi.org:10.1016/j.cell.2007.01.015

13 Hnisz, D. et al. Super-enhancers in the control of cell identity and disease. Cell 155, 934–947 (2013). https://doi.org:10.1016/j.cell.2013.09.053

14 Loven, J. et al. Selective inhibition of tumor oncogenes by disruption of super-enhancers. Cell 153, 320–334 (2013). https://doi.org:10.1016/j.cell.2013.03.036

15 Calo, E. & Wysocka, J. Modification of enhancer chromatin: what, how, and why? Molecular cell 49, 825–837 (2013). https://doi.org:10.1016/j.molcel.2013.01.038

16 Schick, S. et al. Dynamics of chromatin accessibility and epigenetic state in response to UV damage. J Cell Sci 128, 4380–4394 (2015). https://doi.org:10.1242/jcs.173633

17 Guo, L. et al. A combination strategy targeting enhancer plasticity exerts synergistic lethality against BETi-resistant leukemia cells. Nat Commun 11, 740 (2020). https://doi.org:10.1038/s41467-020-14604-6

18 Malik, A. N. et al. Genome-wide identification and characterization of functional neuronal activity-dependent enhancers. Nat Neurosci 17, 1330–1339 (2014). https://doi.org:10.1038/nn.3808

19 Buratowski, S. Progression through the RNA polymerase II CTD cycle. Molecular cell 36, 541–546 (2009). https://doi.org:10.1016/j.molcel.2009.10.019

20 Harlen, K. M. & Churchman, L. S. The code and beyond: transcription regulation by the RNA polymerase II carboxy-terminal domain. Nature reviews. Molecular cell biology 18, 263–273 (2017). https://doi.org:10.1038/nrm.2017.10

21 Chen, F. X., Smith, E. R. & Shilatifard, A. Born to run: control of transcription elongation by RNA polymerase II. Nature reviews. Molecular cell biology 19, 464–478 (2018). https://doi.org:10.1038/s41580-018-0010-5

22 Roy, R. et al. The MO15 cell cycle kinase is associated with the TFIIH transcription-DNA repair factor. Cell 79, 1093–1101 (1994). https://doi.org:10.1016/0092-8674(94)90039-6

23 Jonkers, I. & Lis, J. T. Getting up to speed with transcription elongation by RNA polymerase II. Nat Rev Mol Cell Biol 16, 167–177 (2015). https://doi.org:10.1038/nrm3953

24 Ni, Z. et al. P-TEFb is critical for the maturation of RNA polymerase II into productive elongation in vivo. Molecular and cellular biology 28, 1161–1170 (2008). https://doi.org:10.1128/mcb.01859-07

25 Zheng, B. et al. Acute perturbation strategies in interrogating RNA polymerase II elongation factor function in gene expression. Genes & development 35, 273–285 (2021). https://doi.org:10.1101/gad.346106.120

26 Bugai, A. et al. P-TEFb Activation by RBM7 Shapes a Pro-survival Transcriptional Response to Genotoxic Stress. Molecular cell 74, 254-267.e210 (2019). https://doi.org:10.1016/j.molcel.2019.01.033

27 Ni, Z., Schwartz, B. E., Werner, J., Suarez, J. R. & Lis, J. T. Coordination of transcription, RNA processing, and surveillance by P-TEFb kinase on heat shock genes. Molecular cell 13, 55–65 (2004). https://doi.org:10.1016/s1097-2765(03)00526-4

28 Muse, G. W. et al. RNA polymerase is poised for activation across the genome. Nat Genet 39, 1507–1511 (2007). https://doi.org:10.1038/ng.2007.21

29 Lagha, M. et al. Paused Pol II coordinates tissue morphogenesis in the Drosophila embryo. Cell 153, 976–987 (2013). https://doi.org:10.1016/j.cell.2013.04.045

30 Lin, C. et al. AFF4, a component of the ELL/P-TEFb elongation complex and a shared subunit of MLL chimeras, can link transcription elongation to leukemia. Mol Cell 37, 429–437 (2010). https://doi.org:10.1016/j.molcel.2010.01.026

31 Miller, T. E. et al. Transcription elongation factors represent in vivo cancer dependencies in glioblastoma. Nature 547, 355–359 (2017). https://doi.org:10.1038/nature23000

32 Izumi, K. et al. Germline gain-of-function mutations in AFF4 cause a developmental syndrome functionally linking the super elongation complex and cohesin. Nature genetics 47, 338–344 (2015). https://doi.org:10.1038/ng.3229

33 Dahl, N. A. et al. Super Elongation Complex as a Targetable Dependency in Diffuse Midline Glioma. Cell Rep 31, 107485 (2020). https://doi.org:10.1016/j.celrep.2020.03.049

34 Dahl, N. A. & Vibhakar, R. in Neuro Oncol Vol. 23 1225–1227 (2021).

35 Han, H. et al. TRRUST: a reference database of human transcriptional regulatory interactions. Sci Rep 5, 11432 (2015). https://doi.org:10.1038/srep11432

36 Han, H. et al. TRRUST v2: an expanded reference database of human and mouse transcriptional regulatory interactions. Nucleic Acids Res 46, D380–d386 (2018). https://doi.org:10.1093/nar/gkx1013

37 Muñoz, M. J. et al. DNA damage regulates alternative splicing through inhibition of RNA polymerase II elongation. Cell 137, 708–720 (2009). https://doi.org:10.1016/j.cell.2009.03.010

38 Gao, Y. et al. Acetylation of histone H3K27 signals the transcriptional elongation for estrogen receptor alpha. Commun Biol 3, 165 (2020). https://doi.org:10.1038/s42003-020-0898-0

39 Zhang, Z. et al. Crosstalk between histone modifications indicates that inhibition of arginine methyltransferase CARM1 activity reverses HIV latency. Nucleic Acids Res 45, 9348–9360 (2017). https://doi.org:10.1093/nar/gkx550

40 Hsu, S. C. & Blobel, G. A. The Role of Bromodomain and Extraterminal Motif (BET) Proteins in Chromatin Structure. Cold Spring Harb Symp Quant Biol 82, 37–43 (2017). https://doi.org:10.1101/sqb.2017.82.033829

41 Zhou, Q. & Yik, J. H. The Yin and Yang of P-TEFb regulation: implications for human immunodeficiency virus gene expression and global control of cell growth and differentiation. Microbiol Mol Biol Rev 70, 646–659 (2006). https://doi.org:10.1128/mmbr.00011-06

42 Zhou, Q., Li, T. & Price, D. H. RNA polymerase II elongation control. Annu Rev Biochem 81, 119–143 (2012). https://doi.org:10.1146/annurev-biochem-052610-095910

43 Fortuny, A. & Polo, S. E. The response to DNA damage in heterochromatin domains. Chromosoma 127, 291–300 (2018). https://doi.org:10.1007/s00412-018-0669-6

44 Takata, H. et al. Chromatin compaction protects genomic DNA from radiation damage. PLoS One 8, e75622 (2013). https://doi.org:10.1371/journal.pone.0075622

45 Cidado, J. et al. AZD4573 Is a Highly Selective CDK9 Inhibitor That Suppresses MCL-1 and Induces Apoptosis in Hematologic Cancer Cells. Clinical cancer research : an official journal of the American Association for Cancer Research 26, 922–934 (2020). https://doi.org:10.1158/1078-0432.ccr-19-1853

46 Castel, D. et al. Histone H3F3A and HIST1H3B K27M mutations define two subgroups of diffuse intrinsic pontine gliomas with different prognosis and phenotypes. Acta neuropathologica 130, 815–827 (2015). https://doi.org:10.1007/s00401-015-1478-0

47 Werbrouck, C. et al. TP53 Pathway Alterations Drive Radioresistance in Diffuse Intrinsic Pontine Gliomas (DIPG). Clinical cancer research : an official journal of the American Association for Cancer Research 25, 6788–6800 (2019). https://doi.org:10.1158/1078-0432.ccr-19-0126

48 Metselaar, D. S., du Chatinier, A., Stuiver, I., Kaspers, G. J. L. & Hulleman, E. Radiosensitization in Pediatric High-Grade Glioma: Targets, Resistance and Developments. Frontiers in oncology 11, 662209 (2021). https://doi.org:10.3389/fonc.2021.662209

49 Podhorecka, M., Skladanowski, A. & Bozko, P. H2AX Phosphorylation: Its Role in DNA Damage Response and Cancer Therapy. J Nucleic Acids 2010 (2010). https://doi.org:10.4061/2010/920161

50 Iliakis, G., Wang, Y., Guan, J. & Wang, H. DNA damage checkpoint control in cells exposed to ionizing radiation. Oncogene 22, 5834–5847 (2003). https://doi.org:10.1038/sj.onc.1206682

51 Singh, S. K. et al. Identification of human brain tumour initiating cells. Nature 432, 396–401 (2004). https://doi.org:10.1038/nature03128

52 Chen, J. et al. A restricted cell population propagates glioblastoma growth after chemotherapy. Nature 488, 522–526 (2012). https://doi.org:10.1038/nature11287

53 Bao, S. et al. Glioma stem cells promote radioresistance by preferential activation of the DNA damage response. Nature 444, 756–760 (2006). https://doi.org:10.1038/nature05236

54 Choi, S. A. et al. Identification of brain tumour initiating cells using the stem cell marker aldehyde dehydrogenase. European journal of cancer (Oxford, England : 1990) 50, 137–149 (2014). https://doi.org:10.1016/j.ejca.2013.09.004

55 Surowiec, R. K. et al. Transcriptomic Analysis of Diffuse Intrinsic Pontine Glioma (DIPG) Identifies a Targetable ALDH-Positive Subset of Highly Tumorigenic Cancer Stem-like Cells. Mol Cancer Res 19, 223–239 (2021). https://doi.org:10.1158/1541-7786.mcr-20-0464

56 Wu, G. et al. The genomic landscape of diffuse intrinsic pontine glioma and pediatric non-brainstem high-grade glioma. Nat Genet 46, 444–450 (2014). https://doi.org:10.1038/ng.2938

57 Jones, C. & Baker, S. J. Unique genetic and epigenetic mechanisms driving paediatric diffuse high-grade glioma. Nat Rev Cancer 14 (2014). https://doi.org:10.1038/nrc3811

58 Mackay, A. et al. Integrated Molecular Meta-Analysis of 1,000 Pediatric High-Grade and Diffuse Intrinsic Pontine Glioma. Cancer cell 32, 520-537.e525 (2017). https://doi.org:10.1016/j.ccell.2017.08.017

59 Chang, L., Ruiz, P., Ito, T. & Sellers, W. R. Targeting pan-essential genes in cancer: Challenges and opportunities. Cancer cell 39, 466–479 (2021). https://doi.org:10.1016/j.ccell.2020.12.008

60 Hart, T. et al. High-Resolution CRISPR Screens Reveal Fitness Genes and Genotype-Specific Cancer Liabilities. Cell 163, 1515–1526 (2015). https://doi.org:10.1016/j.cell.2015.11.015

61 Dempster, J. M. et al. Agreement between two large pan-cancer CRISPR-Cas9 gene dependency data sets. Nat Commun 10, 5817 (2019). https://doi.org:10.1038/s41467-019-13805-y

62 Bradner, J. E., Hnisz, D. & Young, R. A. Transcriptional Addiction in Cancer. Cell 168, 629–643 (2017). https://doi.org:10.1016/j.cell.2016.12.013

63 Christensen, C. L. et al. Targeting transcriptional addictions in small cell lung cancer with a covalent CDK7 inhibitor. Cancer cell 26, 909–922 (2014). https://doi.org:10.1016/j.ccell.2014.10.019

64 Wang, Y. et al. CDK7-dependent transcriptional addiction in triple-negative breast cancer. Cell 163, 174–186 (2015). https://doi.org:10.1016/j.cell.2015.08.063

65 Zanconato, F. et al. Transcriptional addiction in cancer cells is mediated by YAP/TAZ through BRD4. Nature medicine 24, 1599–1610 (2018). https://doi.org:10.1038/s41591-018-0158-8

66 Sengupta, S. & George, R. E. Super-Enhancer-Driven Transcriptional Dependencies in Cancer. Trends Cancer 3, 269–281 (2017). https://doi.org:10.1016/j.trecan.2017.03.006

67 Galbraith, M. D., Bender, H. & Espinosa, J. M. Therapeutic targeting of transcriptional cyclin-dependent kinases. Transcription 10, 118–136 (2019). https://doi.org:10.1080/21541264.2018.1539615

68 Gong, L. et al. Differential radiation response between normal astrocytes and glioma cells revealed by comparative transcriptome analysis. Onco Targets Ther 10, 5755–5764 (2017). https://doi.org:10.2147/ott.s144002

69 Baskar, R., Dai, J., Wenlong, N., Yeo, R. & Yeoh, K. W. Biological response of cancer cells to radiation treatment. Front Mol Biosci 1, 24 (2014). https://doi.org:10.3389/fmolb.2014.00024

70 Jones, T. S. & Holland, E. C. Standard of care therapy for malignant glioma and its effect on tumor and stromal cells. Oncogene 31, 1995–2006 (2012). https://doi.org:10.1038/onc.2011.398

71 Liu, X. ABC Family Transporters. Adv Exp Med Biol 1141, 13–100 (2019). https://doi.org:10.1007/978-981-13-7647-4_2

72 Agarwal, S., Hartz, A. M., Elmquist, W. F. & Bauer, B. Breast cancer resistance protein and P-glycoprotein in brain cancer: two gatekeepers team up. Curr Pharm Des 17, 2793–2802 (2011). https://doi.org:10.2174/138161211797440186

73 Le Rhun, E. et al. Profound, durable and MGMT-independent sensitivity of glioblastoma cells to cyclin-dependent kinase inhibition. International journal of cancer 145, 242–253 (2019). https://doi.org:10.1002/ijc.32069

74 Su, Y. T. et al. Novel Targeting of Transcription and Metabolism in Glioblastoma. Clinical cancer research : an official journal of the American Association for Cancer Research 24, 1124–1137 (2018). https://doi.org:10.1158/1078-0432.ccr-17-2032

75 Koncar, R. F. et al. Identification of Novel RAS Signaling Therapeutic Vulnerabilities in Diffuse Intrinsic Pontine Gliomas. Cancer research 79, 4026–4041 (2019). https://doi.org:10.1158/0008-5472.can-18-3521

76 Wu, J. et al. Phase I Study of Zotiraciclib in Combination with Temozolomide for Patients with Recurrent High-grade Astrocytomas. Clin Cancer Res 27, 3298–3306 (2021). https://doi.org:10.1158/1078-0432.ccr-20-4730

77 Curtin, S. C., Minino, A. M. & Anderson, R. N. Declines in Cancer Death Rates Among Children and Adolescents in the United States, 1999–2014. NCHS Data Brief, 1-8 (2016).

78 Luo, Z. et al. The super elongation complex family of RNA polymerase II elongation factors: gene target specificity and transcriptional output. Mol Cell Biol 32, 2608–2617 (2012). https://doi.org:10.1128/mcb.00182-12

79 Ghosh, S. K., Missra, A. & Gilmour, D. S. Negative elongation factor accelerates the rate at which heat shock genes are shut off by facilitating dissociation of heat shock factor. Molecular and cellular biology 31, 4232–4243 (2011). https://doi.org:10.1128/mcb.05930-11

80 Fuda, N. J., Ardehali, M. B. & Lis, J. T. Defining mechanisms that regulate RNA polymerase II transcription in vivo. Nature 461, 186–192 (2009). https://doi.org:10.1038/nature08449

81 Mahat, D. B., Salamanca, H. H., Duarte, F. M., Danko, C. G. & Lis, J. T. Mammalian Heat Shock Response and Mechanisms Underlying Its Genome-wide Transcriptional Regulation. Molecular cell 62, 63–78 (2016). https://doi.org:10.1016/j.molcel.2016.02.025

82 Peterlin, B. M. & Price, D. H. Controlling the elongation phase of transcription with P-TEFb. Mol Cell 23, 297–305 (2006). https://doi.org:10.1016/j.molcel.2006.06.014

83 Luo, Z., Lin, C. & Shilatifard, A. The super elongation complex (SEC) family in transcriptional control. Nat Rev Mol Cell Biol 13, 543–547 (2012). https://doi.org:10.1038/nrm3417

84 Filippakopoulos, P. & Knapp, S. Targeting bromodomains: epigenetic readers of lysine acetylation. Nat Rev Drug Discov 13, 337–356 (2014). https://doi.org:10.1038/nrd4286

85 Liang, K. et al. Targeting Processive Transcription Elongation via SEC Disruption for MYC-Induced Cancer Therapy. Cell 175, 766-779.e717 (2018). https://doi.org:10.1016/j.cell.2018.09.027

86 Katagi, H. et al. Therapeutic targeting of transcriptional elongation in diffuse intrinsic pontine glioma. Neuro-oncology (2021). https://doi.org:10.1093/neuonc/noab009

87 Morales, F. & Giordano, A. Overview of CDK9 as a target in cancer research. Cell Cycle 15, 519–527 (2016). https://doi.org:10.1080/15384101.2016.1138186

88 Chou, J., Quigley, D. A., Robinson, T. M., Feng, F. Y. & Ashworth, A. Transcription-Associated Cyclin-Dependent Kinases as Targets and Biomarkers for Cancer Therapy. Cancer discovery 10, 351–370 (2020). https://doi.org:10.1158/2159-8290.cd-19-0528

89 Karp, J. E. et al. Randomized phase II study of two schedules of flavopiridol given as timed sequential therapy with cytosine arabinoside and mitoxantrone for adults with newly diagnosed, poor-risk acute myelogenous leukemia. Haematologica 97, 1736–1742 (2012). https://doi.org:10.3324/haematol.2012.062539

90 Bible, K. C. et al. A phase 2 trial of flavopiridol (Alvocidib) and cisplatin in platin-resistant ovarian and primary peritoneal carcinoma: MC0261. Gynecol Oncol 127, 55–62 (2012). https://doi.org:10.1016/j.ygyno.2012.05.030

91 Lanasa, M. C. et al. Final results of EFC6663: a multicenter, international, phase 2 study of alvocidib for patients with fludarabine-refractory chronic lymphocytic leukemia. Leuk Res 39, 495–500 (2015). https://doi.org:10.1016/j.leukres.2015.02.001

92 Mita, M. M. et al. Randomized phase II trial of the cyclin-dependent kinase inhibitor dinaciclib (MK-7965) versus capecitabine in patients with advanced breast cancer. Clin Breast Cancer 14, 169–176 (2014). https://doi.org:10.1016/j.clbc.2013.10.016

93 Stephenson, J. J. et al. Randomized phase 2 study of the cyclin-dependent kinase inhibitor dinaciclib (MK-7965) versus erlotinib in patients with non-small cell lung cancer. Lung Cancer 83, 219–223 (2014). https://doi.org:10.1016/j.lungcan.2013.11.020

94 Gojo, I. et al. Clinical and laboratory studies of the novel cyclin-dependent kinase inhibitor dinaciclib (SCH 727965) in acute leukemias. Cancer chemotherapy and pharmacology 72, 897–908 (2013). https://doi.org:10.1007/s00280-013-2249-z

95 Roboz, G. J. et al. Phase I dose escalation study of TG02 in patients with advanced hematologic malignancies. Journal of Clinical Oncology 30, 6577–6577 (2012). https://doi.org:10.1200/jco.2012.30.15_suppl.6577

96 Balakrishnan, I. et al. Senescence Induced by BMI1 Inhibition Is a Therapeutic Vulnerability in H3K27M-Mutant DIPG. Cell Rep 33, 108286 (2020). https://doi.org:10.1016/j.celrep.2020.108286

97 Hu, Y. & Smyth, G. K. ELDA: extreme limiting dilution analysis for comparing depleted and enriched populations in stem cell and other assays. Journal of immunological methods 347, 70–78 (2009). https://doi.org:10.1016/j.jim.2009.06.008

98 Zhou, Y. et al. Metascape provides a biologist-oriented resource for the analysis of systems-level datasets. Nat Commun 10, 1523 (2019). https://doi.org:10.1038/s41467-019-09234-6

99 Andrews, S. (Babraham Bioinformatics, 2010).

100 Ewels, P., Magnusson, M., Lundin, S. & Käller, M. MultiQC: summarize analysis results for multiple tools and samples in a single report. Bioinformatics 32, 3047–3048 (2016). https://doi.org:10.1093/bioinformatics/btw354

101 Langmead, B. & Salzberg, S. L. Fast gapped-read alignment with Bowtie 2. Nat Methods 9, 357–359 (2012). https://doi.org:10.1038/nmeth.1923

102 Li, H. et al. The Sequence Alignment/Map format and SAMtools. Bioinformatics 25, 2078–2079 (2009). https://doi.org:10.1093/bioinformatics/btp352

103 Quinlan, A. R. & Hall, I. M. BEDTools: a flexible suite of utilities for comparing genomic features. Bioinformatics 26, 841–842 (2010). https://doi.org:10.1093/bioinformatics/btq033

104 Ramírez, F. et al. deepTools2: a next generation web server for deep-sequencing data analysis. Nucleic Acids Res 44, W160–165 (2016). https://doi.org:10.1093/nar/gkw257

105 Thorvaldsdottir, H., Robinson, J. T. & Mesirov, J. P. Integrative Genomics Viewer (IGV): high-performance genomics data visualization and exploration. Brief Bioinform 14, 178–192 (2013). https://doi.org:10.1093/bib/bbs017

106 Zhang, Y. et al. Model-based analysis of ChIP-Seq (MACS). Genome Biol 9, R137 (2008). https://doi.org:10.1186/gb-2008-9-9-r137

107 Li, Q., Brown, J. B., Huang, H. & Bickel, P. J. Measuring reproducibility of high-throughput experiments. The Annals of Applied Statistics 5, 1752-1779, 1728 (2011).

108 Riemondy, K. A. et al. valr: Reproducible genome interval analysis in R. F1000Res 6, 1025 (2017). https://doi.org:10.12688/f1000research.11997.1

109 Stark, R. (ed Gord Brown) (Bioconductor, 2011).

110 Shen, L., Shao, N., Liu, X. & Nestler, E. ngs.plot: Quick mining and visualization of next-generation sequencing data by integrating genomic databases. BMC Genomics 15, 284 (2014). https://doi.org:10.1186/1471-2164-15-284

111 Madhavan, K. et al. Venetoclax cooperates with ionizing radiation to attenuate Diffuse Midline Glioma tumor growth. Clin Cancer Res (2022). https://doi.org:10.1158/1078-0432.ccr-21-4002

