## Supplemental Figures for "Rapid PTEFb-dependent transcriptional reorganization underpins the glioma adaptive response to radiotherapy"

**Extended Data**

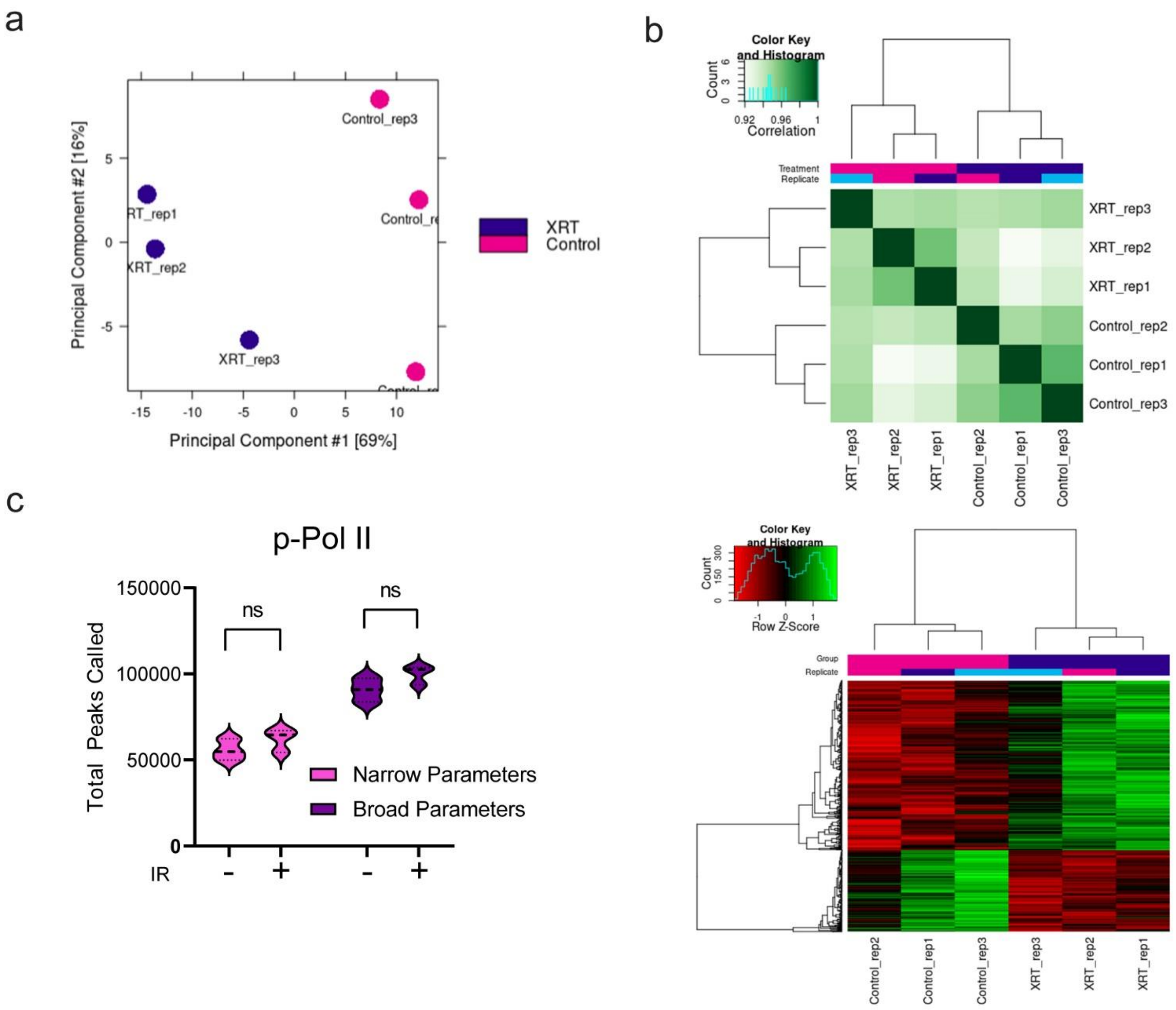


**Extended Data Figure 1.** p-Pol II CUT&RUN reveals distinct occupancy changes following IR-exposure. **a.** Principle component analysis of differentially bound, reproducible peaks. **b.** Unsupervised hierarchical clustering of reproducible p-Pol II peak calls demonstrates concordance within sample replicates. **c.** Total peaks called using MACS2 narrow or broad peak parameters. Quantitative comparisons reflect two-tailed Student’s t-test.

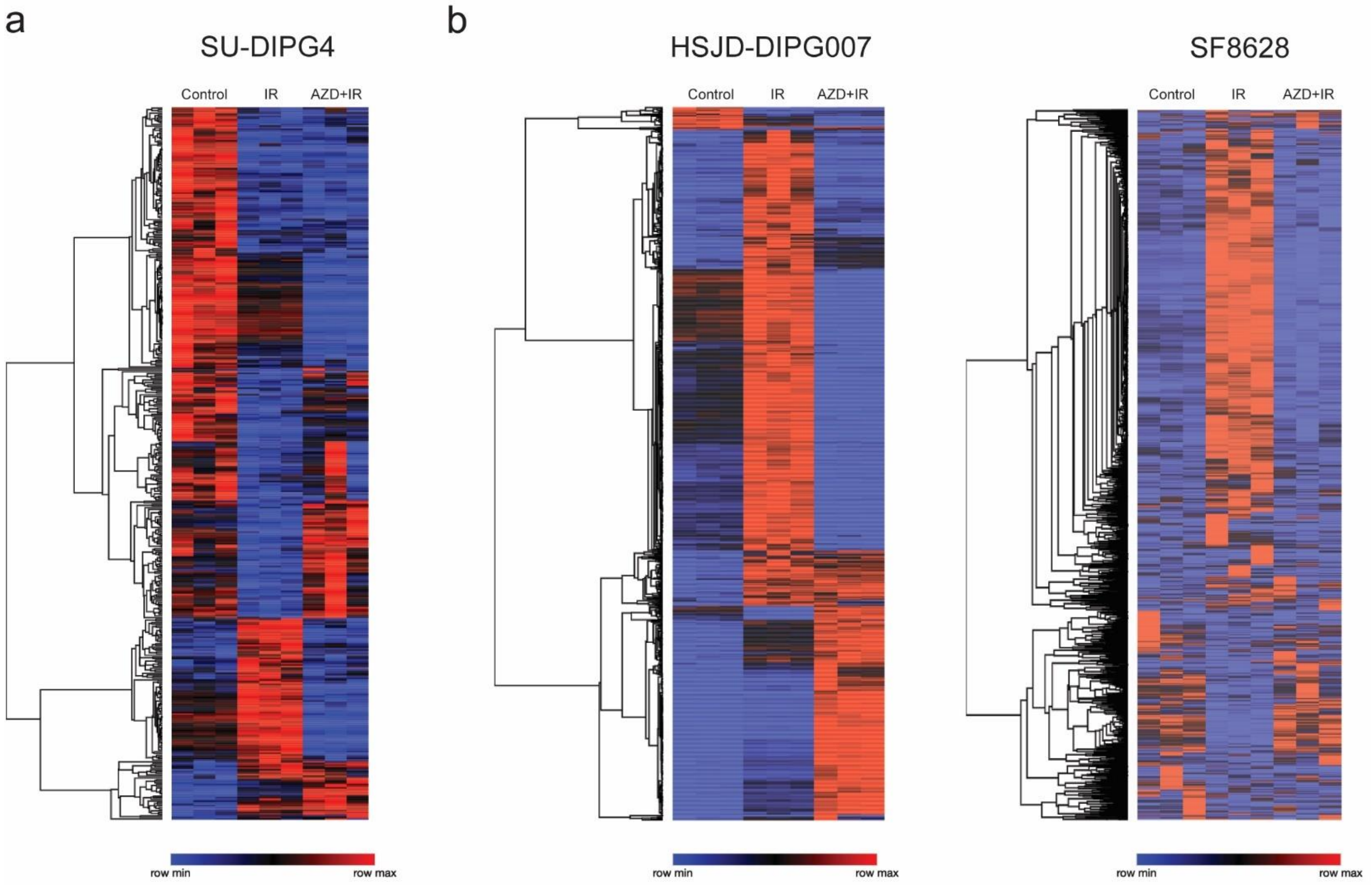


**Extended Data Figure 2.** IR-induced expression profiles of DMG cell lines. **a.** Unsupervised hierarchical clustering of SU-DIPG4 gene expression LFC ≥ +/- 1.2 in control vs IR samples (n=3, padj <0.05). **b.** Unsupervised hierarchical clustering of HSJD-DIPG7 and SF8628 gene expression LFC ≥ +/- 1.2 in control vs IR samples (n=3, padj <0.05).

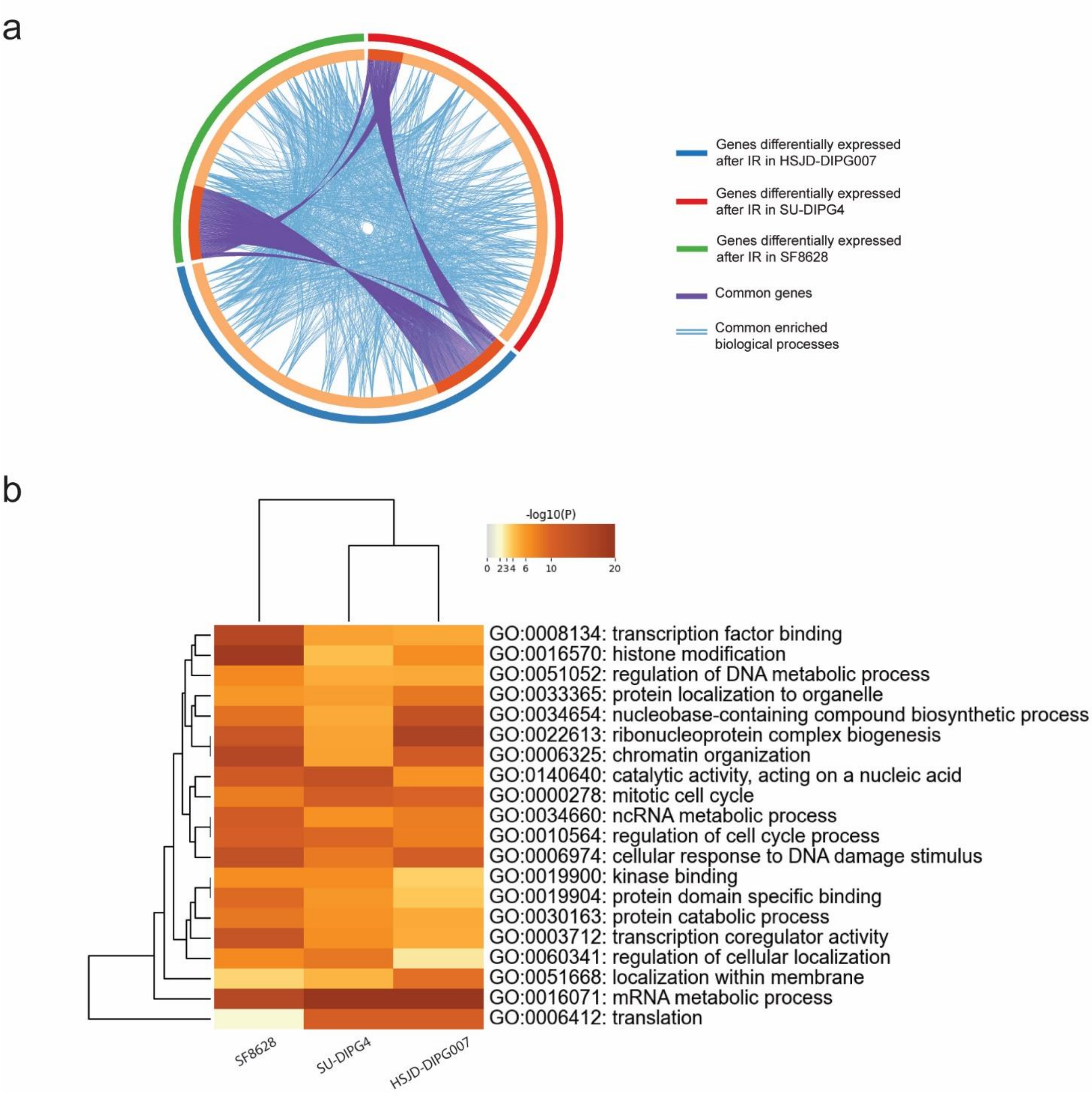


**Extended Data Figure 3.** IR-induced expression changes in pHGG models. **a.** Circos plot of genes upregulated in respective models following single IR exposure. Identical genes are indicated in purple, while common biological processes are indicated in blue. **b.** Unsupervised hierarchical clustering of common ontology terms from (a).

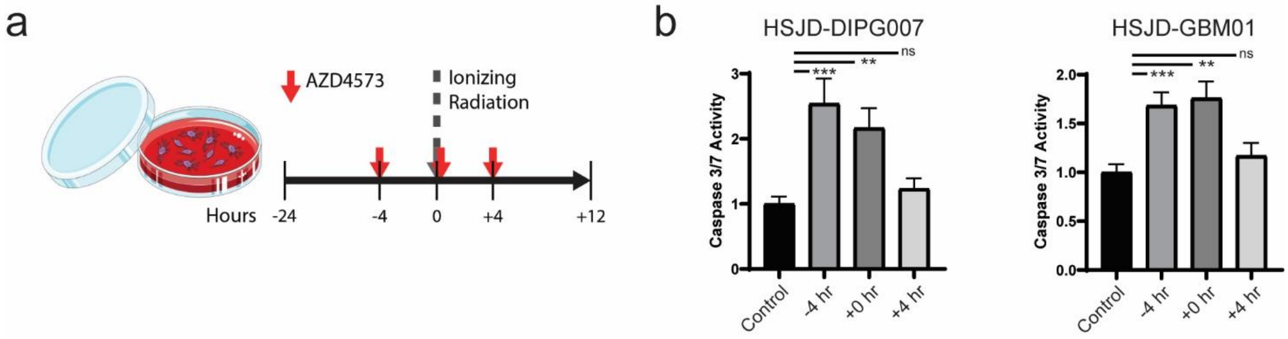


**Extended Data Figure 4.** Sequential optimization of AZD4573 and IR combinatorial therapy. **a.** Schematic of the experiment in which a fixed 6 nM dose of AZD4573 was added at various timepoints relative to single 4 Gy IR treatment. Caspase 3/7 activity was then measured at 12 hours following IR. Longer timepoints did not alter relative ratio of caspase induction (data not shown). **b.** Relative caspase 3/7 activity of HGG cultures treated with IR (control) and AZD4573 at indicated timepoints. Quantitative comparisons reflect two-tailed Student's t-test (** p=<0.01, *** p=<0.001).

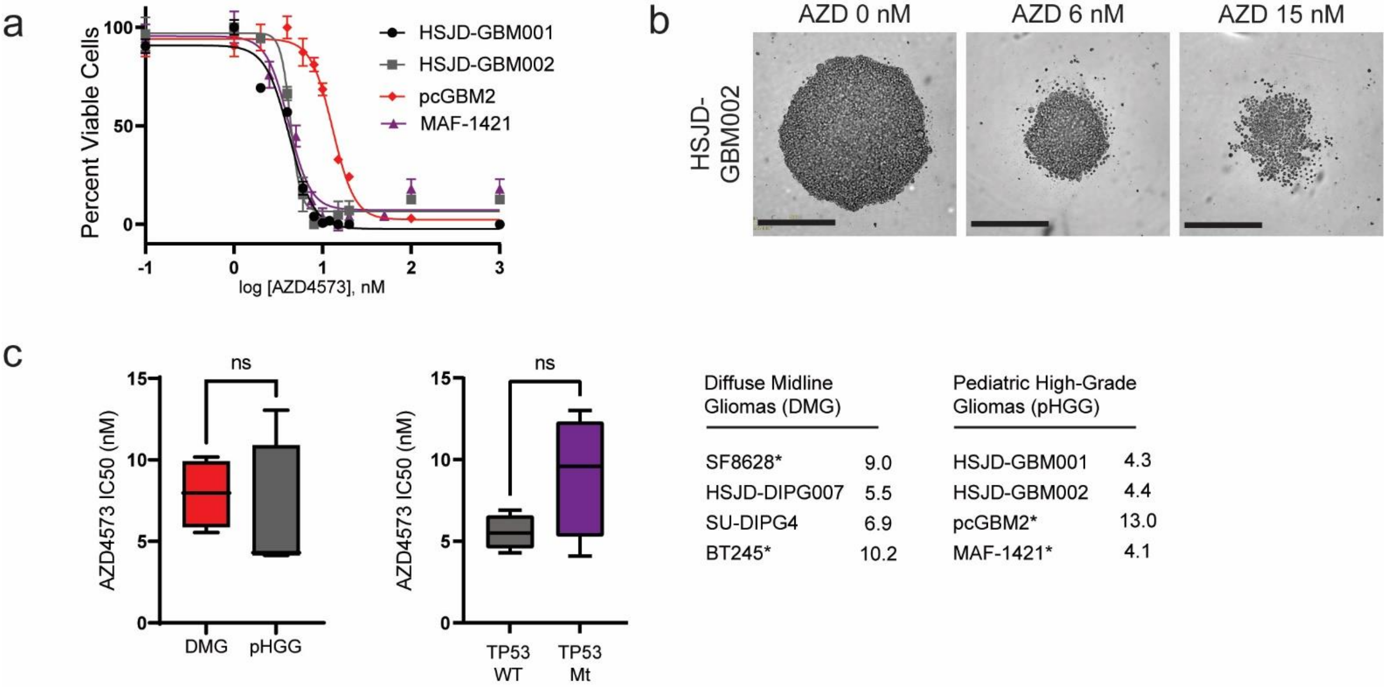


**Extended Data Figure 5.** AZD4573 demonstrates *in vitro* efficacy across the mutational spectrum of pHGG. **a.** Dose response curve of AZD4573 across a panel of H3 WT (HSJD-GBM001, pcGBM2, MAF-1421) and H3G34R mutant (HSJD-GBM002) glioma cultures. Corresponding dose response curves in panel of H3K27M cultures previously published in (Dahl et al., 2020). **b.** Representative live cell imaging of HSJD-GBM002 neurosphere cultures at indicated concentrations of AZD4573 (scale bars, 400 μM). **c.** Mean half-maximal inhibitory concentration of AZD4573 in H3K27M-mutant cultures (DMG) in comparison to H3 WT or H3G34R-mutant (pHGG) (left) or *TP53* wild-type cultures versus *TP53* mutant (center). Quantitative comparison reflects two-tailed Student's t-test (left). Individual culture models and $IC_{50}$ values in nM (right), * indicates *TP53* mutant status.

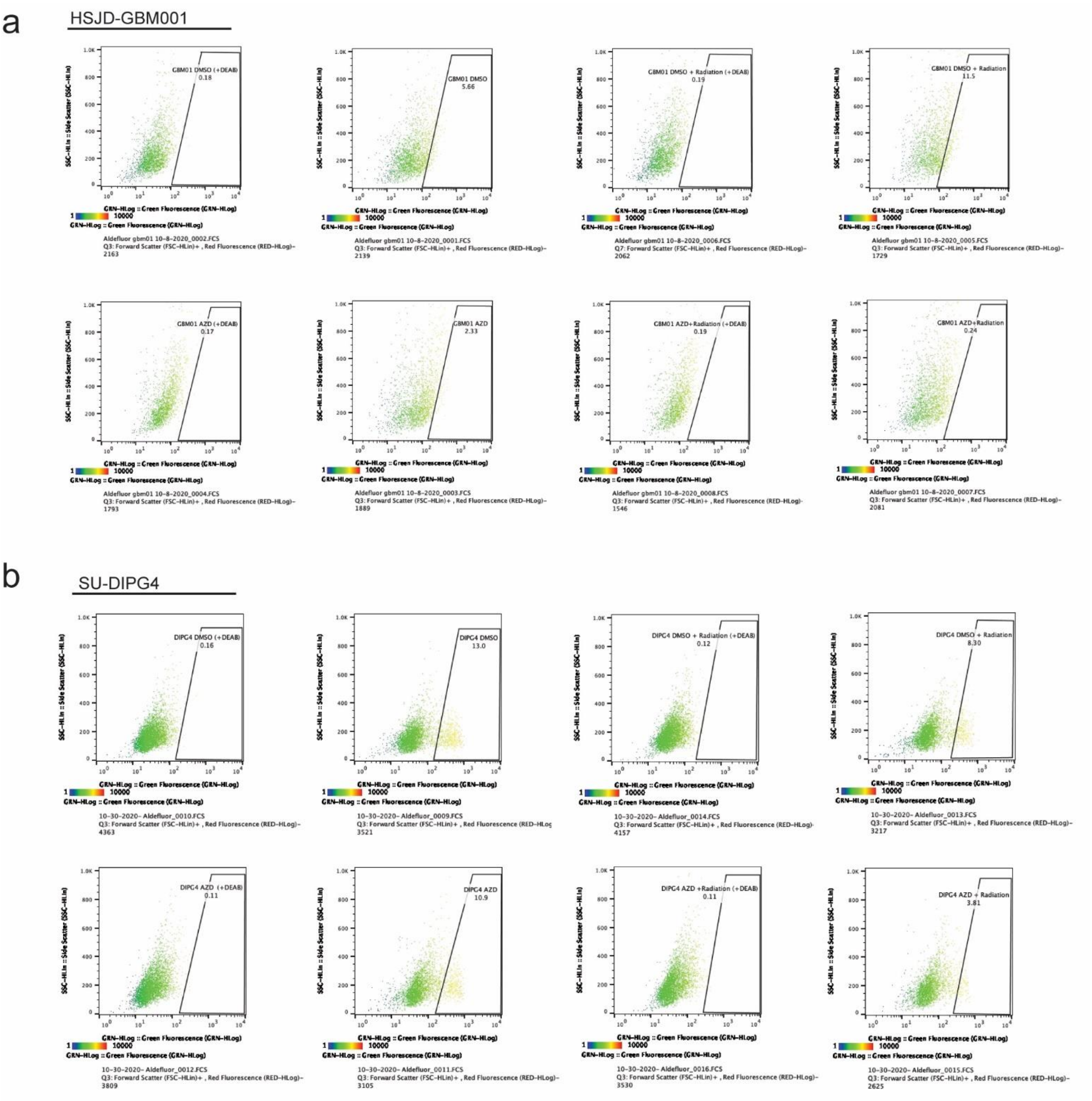


**Extended Data Figure 6.** AZD4573 and IR combinatorial therapy effectively deplete brain tumor initiating cell fraction within culture models. Brain tumor initiating cell fraction after DMSO control, AZD4573 (5nM), IR (4 Gy), or combination treatment as identified by ALDH expression in (**a**) H3 WT pHGG (HSJD-GBM001) and (**b**) H3K27M-mutant DMG (SU-DIPG4).

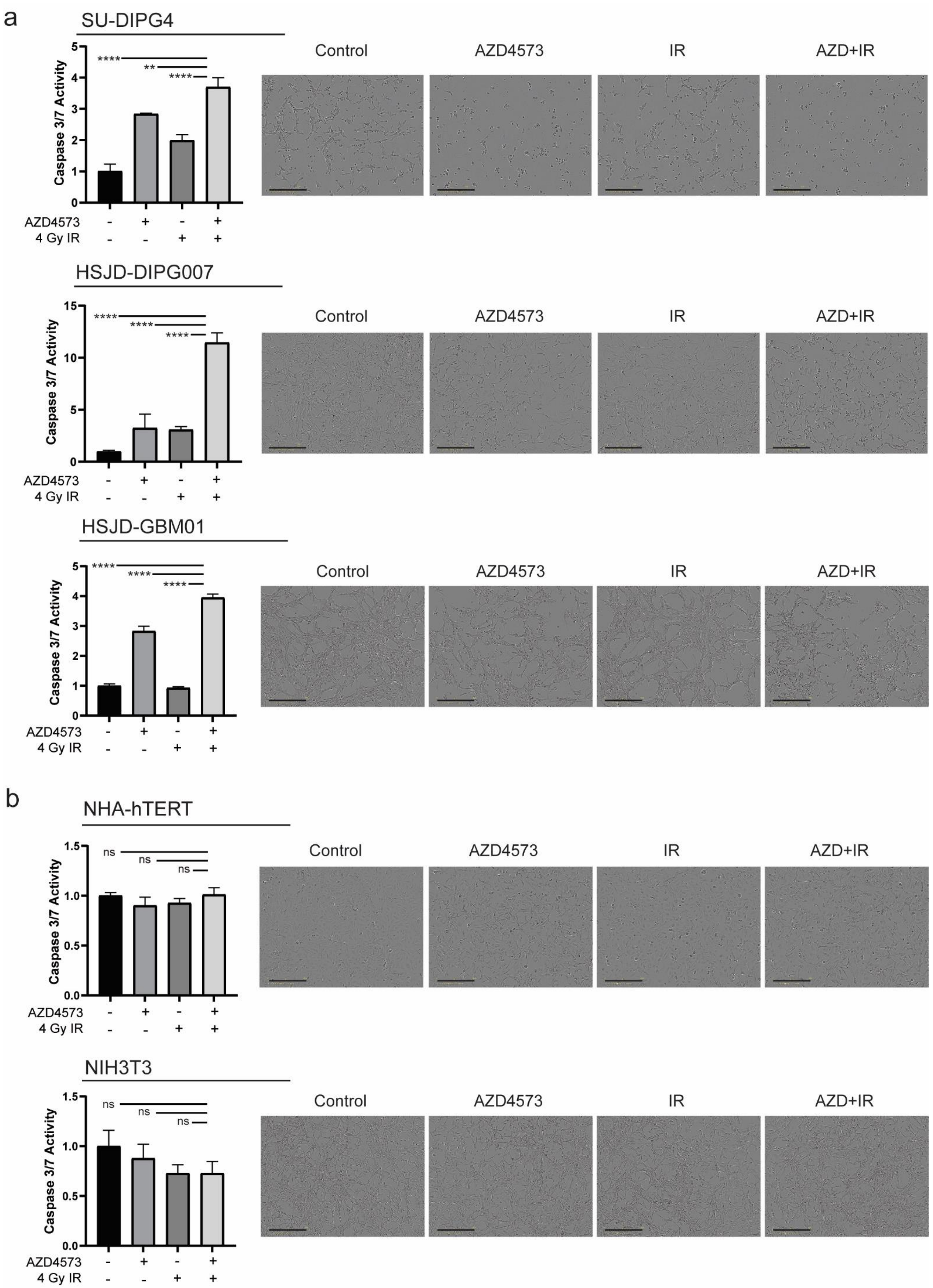

a
SU-DIPG4
Caspase 3/7 Activity
****
**
****
AZD4573
4 Gy IR
Control
AZD4573
IR
AZD+IR
HSJD-DIPG007
Caspase 3/7 Activity
****
****
****
AZD4573
4 Gy IR
Control
AZD4573
IR
AZD+IR
HSJD-GBM01
Caspase 3/7 Activity
****
****
****
AZD4573
4 Gy IR
Control
AZD4573
IR
AZD+IR
b
NHA-hTERT
Caspase 3/7 Activity
ns
ns
ns
AZD4573
4 Gy IR
Control
AZD4573
IR
AZD+IR
NIH3T3
Caspase 3/7 Activity
ns
ns
ns
AZD4573
4 Gy IR
Control
AZD4573
IR
AZD+IR

**Extended Data Figure 7.** Therapeutic index for CDK9i and IR in HGG cultures relative to normal controls. Caspase 3/7 activity (24 hours, left) and representative live-cell imaging (36 hours, right) of pediatric HGG cultures (**a**) or normal cell controls (**b**) after treatment with DMSO control, AZD4573, IR, or combination. Quantitative comparisons reflect two-tailed Student's t-test (** p=<0.01, *** p=<0.001, **** p=<0.0001).

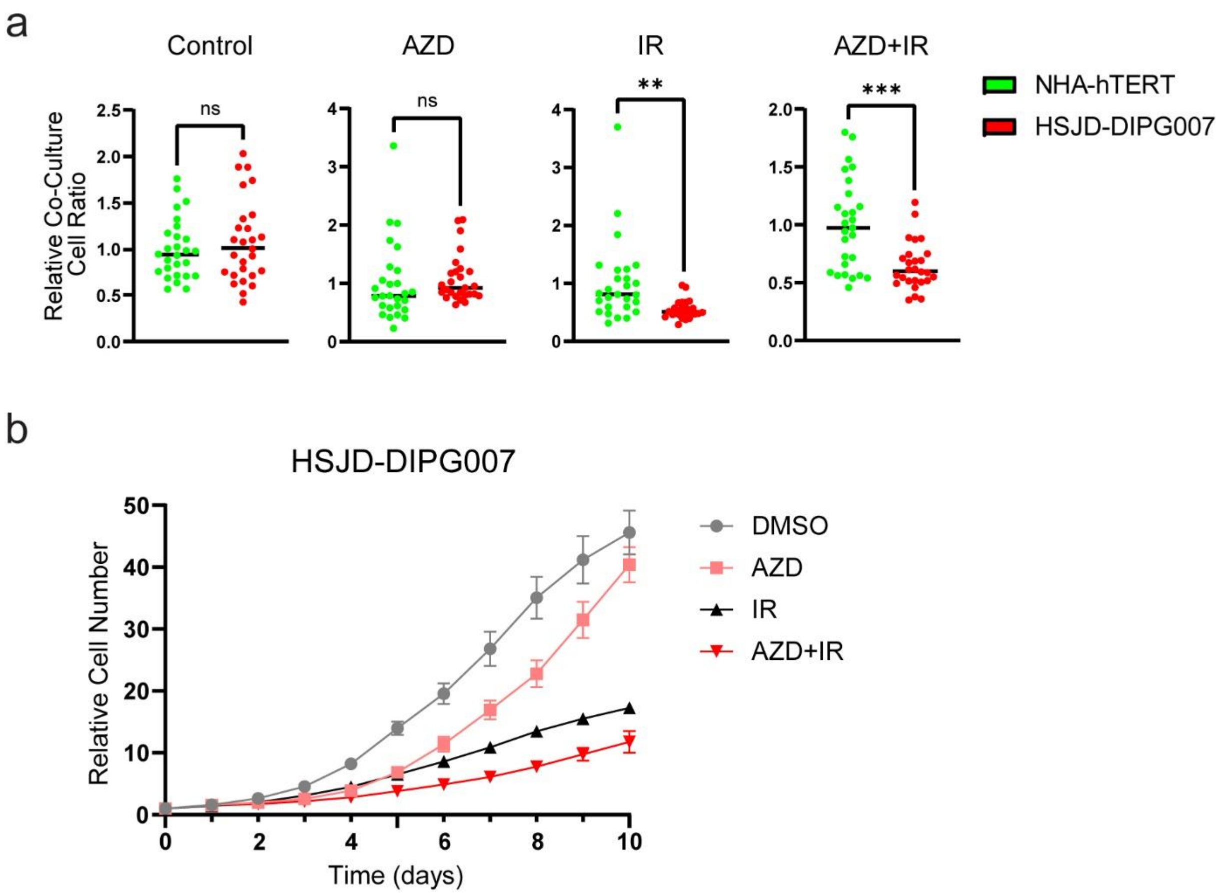


**Extended Data Figure 8.** Therapeutic index of combinatorial therapy within a co-culture system. **a.** Relative ratio quantification of co-cultured DIPG cell and astrocytes at day 10 following indicated treatments. Quantitative comparisons reflect two-tailed Student's t-test (** p=<0.01, *** p=<0.001). **b.** Relative number of HSJD-DIPG7 cells from co-culture system at indicated timepoints, shown as mean ± SEM.

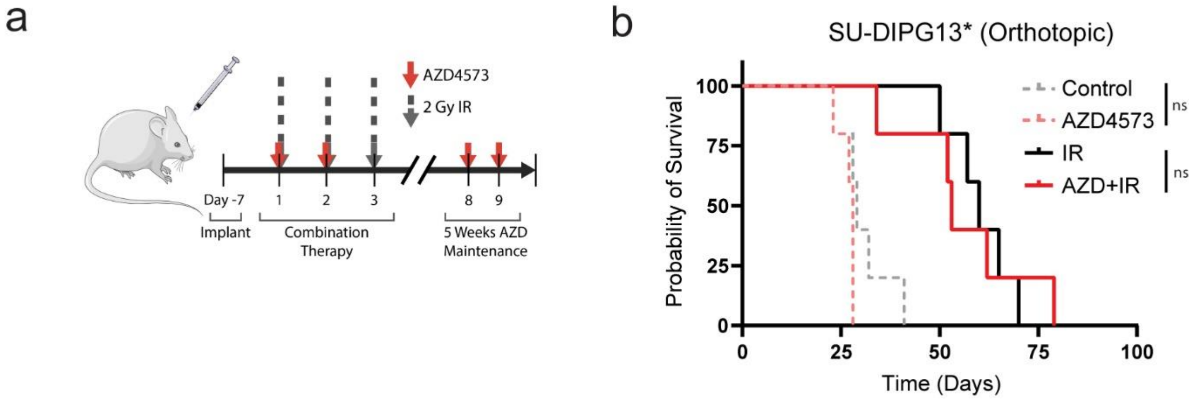


**Extended Data Figure 9.** AZD4573 is ineffective against intracranial model of DIPG. **a.** Schematic represents the treatment schedule of SU-DIPG13* xenografts with either AZD4573 (15/15 mg/kg biweekly administered intraperitoneally), radiotherapy (2 Gy x 3 fractions), or combination. **b.** Kaplan-Meier survival analysis of orthotopic xenograft cohorts from (a) receiving indicated treatments.

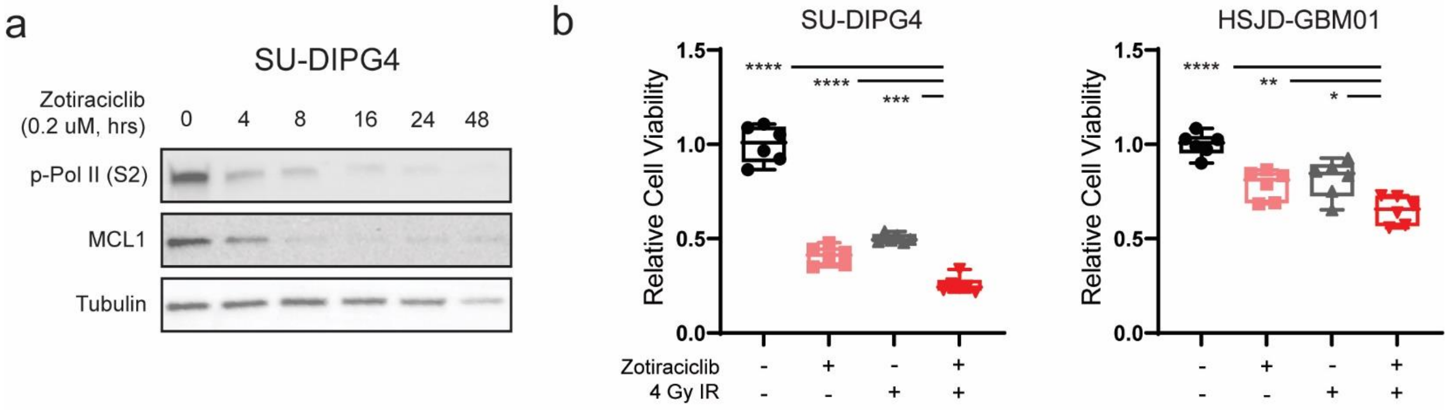


**Extended Data Figure 10.** CDK9 inhibitory activity of zotiraciclib. **a.** Western blot analysis of p-Pol II (Ser 2) and MCL1 after indicated exposure times to 0.2 µM zotiraciclib. **b.** Cell viability measured at 3 days following 24-hour exposure to 0.2 µM zotiraciclib +/- 4 Gy IR. Box whiskers represent min to max range of replicates. Quantitative comparisons reflect two-tailed Student's t-test (* p=<0.05, ** p=<0.01, *** p=<0.001, **** p=<0.0001).

| Cell Line | Marker | Allele 1 | Allele 2 |
|---|---|---|---|
| BT245 | D3S1358 | 18 | |
| BT245 | vWA | 14 | 15 |
| BT245 | D16S539 | 11 | |
| BT245 | CSF1PO | 12 | |
| BT245 | TPOX | 8 | 12 |
| BT245 | Yindel | 2 | |
| BT245 | AMEL | X | Y |
| BT245 | D8S1179 | 11 | 13 |
| BT245 | D21S11 | 31.2 | |
| BT245 | D18S51 | 14 | |
| BT245 | DYS391 | 9 | |
| BT245 | D2S441 | 11.3 | 14 |
| BT245 | D19S433 | 14 | 15 |
| BT245 | TH01 | 9 | |
| BT245 | FGA | 19 | |
| BT245 | D22S1045 | 15 | 16 |
| BT245 | D5S818 | 12 | |
| BT245 | D13S317 | 11 | |
| BT245 | D7S820 | 8 | 10 |
| BT245 | SE33 | 16 | |
| BT245 | D10S1248 | 12 | |
| BT245 | D1S1656 | 16 | 17.3 |
| BT245 | D12S391 | 23 | 24 |
| BT245 | D2S1338 | 17 | 18 |

| Cell Line | Marker | Allele 1 | Allele 2 |
|---|---|---|---|
| GBM01 | D3S1358 | 15 | |
| GBM01 | vWA | 17 | 18 |
| GBM01 | D16S539 | 12 | |
| GBM01 | CSF1PO | 9 | |
| GBM01 | TPOX | 8 | 10 |
| GBM01 | Yindel | | |
| GBM01 | AMEL | X | |
| GBM01 | D8S1179 | 10 | 13 |
| GBM01 | D21S11 | 30 | |
| GBM01 | D18S51 | 18 | |
| GBM01 | DYS391 | | |
| GBM01 | D2S441 | 11.3 | 13 |
| GBM01 | D19S433 | 13 | 14 |
| GBM01 | TH01 | 8 | 9.3 |
| GBM01 | FGA | 25 | |
| GBM01 | D22S1045 | 16 | |
| GBM01 | D5S818 | 13 | 14 |
| GBM01 | D13S317 | 12 | 14 |
| GBM01 | D7S820 | 10 | 11 |
| GBM01 | SE33 | 21 | 28.2 |
| GBM01 | D10S1248 | 13 | 16 |
| GBM01 | D1S1656 | 15 | |
| GBM01 | D12S391 | 18 | 19 |
| GBM01 | D2S1338 | 19 | 20 |

| Cell Line | Marker | Allele 1 | Allele 2 |
|---|---|---|---|
| DIPG4 | D3S1358 | 14 | 16 |
| DIPG4 | vWA | 15 | 19 |
| DIPG4 | D16S539 | 9 | |
| DIPG4 | CSF1PO | 9 | 10 |
| DIPG4 | TPOX | 8 | |
| DIPG4 | Yindel | | |
| DIPG4 | AMEL | X | |
| DIPG4 | D8S1179 | 10 | 12 |
| DIPG4 | D21S11 | 29 | 31 |
| DIPG4 | D18S51 | 14 | |
| DIPG4 | DYS391 | | |
| DIPG4 | D2S441 | 11 | |
| DIPG4 | D19S433 | 13 | 14 |
| DIPG4 | TH01 | 6 | 9.3 |
| DIPG4 | FGA | 24 | |
| DIPG4 | D22S1045 | 15 | |
| DIPG4 | D5S818 | 12 | 13 |
| DIPG4 | D13S317 | 7 | 12 |
| DIPG4 | D7S820 | 10 | 11 |
| DIPG4 | SE33 | 24.2 | 28.2 |
| DIPG4 | D10S1248 | 15 | 16 |
| DIPG4 | D1S1656 | 17 | 18.3 |
| DIPG4 | D12S391 | 22 | 25 |
| DIPG4 | D2S1338 | 20 | 24 |

| Cell Line | Marker | Allele 1 | Allele 2 |
|---|---|---|---|
| DIPG7 | D3S1358 | 16 | 17 |
| DIPG7 | vWA | 16 | 17 |
| DIPG7 | D16S539 | 13 | 14 |
| DIPG7 | CSF1PO | 10 | |
| DIPG7 | TPOX | 8 | |
| DIPG7 | Yindel | | |
| DIPG7 | AMEL | X | Y |
| DIPG7 | D8S1179 | 11 | 13 |
| DIPG7 | D21S11 | 29 | 30 |
| DIPG7 | D18S51 | 19 | |
| DIPG7 | DYS391 | | |
| DIPG7 | D2S441 | 11 | |
| DIPG7 | D19S433 | 12 | 14 |
| DIPG7 | TH01 | 6 | 9.3 |
| DIPG7 | FGA | 21 | 23 |
| DIPG7 | D22S1045 | 16 | |
| DIPG7 | D5S818 | 12 | 13 |
| DIPG7 | D13S317 | 13 | |
| DIPG7 | D7S820 | 9 | 12 |
| DIPG7 | SE33 | 21.2 | |
| DIPG7 | D10S1248 | 12 | 14 |
| DIPG7 | D1S1656 | 15 | 16 |
| DIPG7 | D12S391 | 17 | 20 |
| DIPG7 | D2S1338 | 23 | 24 |

| Cell Line | Marker | Allele 1 | Allele 2 |
|---|---|---|---|
| DIPG13 | D3S1358 | 17 | |
| DIPG13 | vWA | 18 | 19 |
| DIPG13 | D16S539 | 11 | 12 |
| DIPG13 | CSF1PO | 9 | 10 |
| DIPG13 | TPOX | 8 | 9 |
| DIPG13 | Yindel | | |
| DIPG13 | AMEL | X | |
| DIPG13 | D8S1179 | 9 | 14 |
| DIPG13 | D21S11 | 30 | 31 |
| DIPG13 | D18S51 | 14 | 18 |
| DIPG13 | DYS391 | | |
| DIPG13 | D2S441 | 11 | |
| DIPG13 | D19S433 | 12 | 13 |
| DIPG13 | TH01 | 6 | |
| DIPG13 | FGA | 20 | 22 |
| DIPG13 | D22S1045 | 11 | 16 |
| DIPG13 | D5S818 | 12 | |
| DIPG13 | D13S317 | 11 | |
| DIPG13 | D7S820 | 9 | |
| DIPG13 | SE33 | 23.2 | 29.2 |
| DIPG13 | D10S1248 | 13 | 14 |
| DIPG13 | D1S1656 | 12 | |
| DIPG13 | D12S391 | 16 | 17 |
| DIPG13 | D2S1338 | 23 | |

| Cell Line | Marker | Allele 1 | Allele 2 |
|---|---|---|---|
| SF8628 | D3S1358 | 15 | 18 |
| SF8628 | vWA | 16 | 17 |
| SF8628 | D16S539 | 9 | |
| SF8628 | CSF1PO | 11 | 12 |
| SF8628 | TPOX | 7.3 | |
| SF8628 | Yindel | | |
| SF8628 | AMEL | X | |
| SF8628 | D8S1179 | 10 | 13 |
| SF8628 | D21S11 | 29 | 30 |
| SF8628 | D18S51 | 14 | |
| SF8628 | DYS391 | | |
| SF8628 | D2S441 | 10 | 11 |
| SF8628 | D19S433 | 12 | 14 |
| SF8628 | TH01 | 7 | |
| SF8628 | FGA | 19 | 22 |
| SF8628 | D22S1045 | 15 | 16 |
| SF8628 | D5S818 | 11 | 12 |
| SF8628 | D13S317 | 9 | |
| SF8628 | D7S820 | 11 | |
| SF8628 | SE33 | 27.2 | 28.2 |
| SF8628 | D10S1248 | 13 | 15 |
| SF8628 | D1S1656 | 16 | 17 |
| SF8628 | D12S391 | 19 | 21 |
| SF8628 | D2S1338 | 17 | 18 |

| Cell Line | Marker | Allele 1 | Allele 2 |
|---|---|---|---|
| NHA | D3S1358 | 15 | 16 |
| NHA | vWA | 18 | |
| NHA | D16S539 | 12 | |
| NHA | CSF1PO | 11 | 12 |
| NHA | TPOX | 8 | |
| NHA | Yindel | | |
| NHA | AMEL | X | |
| NHA | D8S1179 | 13 | 14 |
| NHA | D21S11 | 31 | 32.2 |
| NHA | D18S51 | 14 | 17 |
| NHA | DYS391 | | |
| NHA | D2S441 | 10 | 11 |
| NHA | D19S433 | 13 | 15 |
| NHA | TH01 | 6 | 8 |
| NHA | FGA | 21 | 23 |
| NHA | D22S1045 | 11 | 16 |
| NHA | D5S818 | 11 | |
| NHA | D13S317 | 8 | 11 |
| NHA | D7S820 | 8 | 11 |
| NHA | SE33 | 19 | 31.2 |
| NHA | D10S1248 | 13 | 15 |
| NHA | D1S1656 | 12 | 17.3 |
| NHA | D12S391 | 20 | 23 |
| NHA | D2S1338 | 17 | 19 |

**Extended Data Figure 11.** STR profiling of cell culture models.
